# New genes play a prominent role in evolution of new sperm classes of *Drosophila*

**DOI:** 10.64898/2026.07.10.737682

**Authors:** Huangyi He, Qianwei Su, Lu Cao, Mujahid Ali, Lubna Younas, Junbo Jia, Mingyue Wang, Rong Xiang, Xun Xu, Luonan Chen, Qi Zhou

## Abstract

The widespread and regulated production of infertile parasperm assisting fertilization of eusperm provides a compelling model for elucidating the mechanisms of cellular innovation and altruism. Here we identify two specific parasperm lineages in *Drosophila pseudoobscura* that are segregated in development from the eusperm lineage by their single-cell transcriptomes. Most X-linked genes, except for species-specific new genes, show testis-specific repressive epigenetic modifications and become downregulated upon meiotic entry, supporting meiotic X chromosome inactivation. Between sperm classes, nearly half of the known ‘meiotic arrest’ genes and their downstream targets are differentially transcribed, with over 200 genes displaying a gradient of transcription levels correlated with that of tail lengths. New genes are overrepresented throughout the hierarchical regulatory network of sperm divergence and over 60% of them exhibit a new cell-level expression compared to their parental copies. These results together indicate that new genes play a prominent role in the emergence of low-cost parasperm that shield the eusperm until fertilization.

## Introduction

Testes are one of the fastest-evolving organs despite their ancient origin, due to the intense sexual selection^1^ and unique transcriptional regulation. Compared to somatic organs, including the brain with many more cell types, testis has a more widespread and complex transcriptome. In both mammalian and *Drosophila* testes, the majority of protein-coding genes and otherwise silenced intergenic regions in the soma are transcribed^2–4^, possibly due to the transcription coupled DNA repair or chromatin remodeling during spermatogenesis^5,6^. The permissive transcription environment, together with meiotic sex chromosome inactivation (MSCI) in testis have profound and complex impacts on genome evolution. In particular, these forces influence the origin of new genes and their non-random chromosomal distribution. Testes have been hypothesized to acts as a ‘cradle for new genes’ due to species specific genes, including *de novo* genes originated from non-coding sequences and new duplicated genes from pre-existing parental genes, often have biased expressions in the male germline^2,7^. New genes may subsequently become fixed in the population by haploid^8^ and sexual selection^9^. In particular, population genetic models predict the hemizygous X chromosome is a hotspot for nascent genes, since it can more easily fix recessive beneficial alleles than autosomes^10^. On the other hand, X chromosomes of many animal species exhibit a pronounced deficiency of testis genes (so-called ‘demasculinization’), either as a disfavored chromosomal location for male meiotic genes due to MSCI^11^, and/or because it is more often transmitted in females compared to autosomes^12^.

An important hypothesis with only emerging experimental evidence is that new genes may originate as the genome substrate for the innovation of cell types in specific lineages^13–16^. Testis, where new genes are preferentially transcribed, comprise a potent model for testing this hypothesis because sperm have evolved the greatest variation of morphology among cell types^17,18^, including novel species-specific morphs that sometimes co-exist with the ancestral cell morph. It is estimated that half of the examined invertebrate (e.g., some molluscs, insects, nematodes etc.) taxonomical groups and a few vertebrate (e.g., some fish) species independently evolved sperm heteromorphism, which refers to simultaneous production of one class of fertile eusperm (or called eupyrene in Lepidoptera), and one or more classes of infertile parasperm (or apyrene) ^19^. Such ubiquitous sperm heteromorphism is not a product of meiotic errors, because a stable and large (e.g., 34%-94% of the ejaculate among *Drosophila* species^20^) proportion of parasperm is observed, suggesting a tightly regulated but uncharacterized developmental program evolved specifically in those species. A broader question is that, why and how a new morph/state of cells, like parasperm, would sacrifice its primary function to fulfill that of other cells?

Therefore, species that evolved sperm heteromorphism offer a paradigm for understanding the molecular and evolutionary mechanisms of cell-level novelty and altruism^21^. One of the best characterized species to address this fundamental question is *D. pseudoobscura* (*Dpse*). Compared to other *obscura* group species with one eusperm (Eu) and one parasperm class, *Dpse* and its sister species *D. persimilis* and *D. miranda* produce an additional parasperm class (Para1 and Para2) ^22^. The tails of three classes form a length gradient but their ultrastructural features are indistinguishable (**Fig. 1a, and Supplementary Fig. 1-2**), similar to reported results in *D. subobscura*^23^. Several mutually non-exclusive hypotheses have been proposed to account for the functions of parasperm. They are assumed to often evolve with a lower energy cost to protect eusperm as a ‘cheap shield’ from spermicide or against the harsh and cryptic selection by the female reproductive tract^24–27^, or to provide nutrients to the eusperm or female^19,28^, or to assist eusperm to compete with those of other male individuals (e.g., by blocking them or filling up the female reproductive tract, thereby acting as ‘cheap fillers’) ^29^. In *Dpse*, the cheap shield hypothesis is supported by the positive correlation between the proportion of parasperm and the eusperm viability, when the ejaculate is exposed to the extract of female reproductive tract^26,27^. The cheap-filler and nutrient hypotheses of parasperm have not been supported^20,30,31^. Para2 with the medium tail length was recently reported to have a distinct corkscrew tail shape that was implicated to be associated with sperm competition, with an unknown mechanism^26^. Another distinction of *Dpse* from most other obscura species is that it has experienced complex turnovers of sex chromosomes: the ancestral *Drosophila* X chromosome (termed the ‘Muller-A’ element across *Drosophila* species, AX) has fused to one autosome (‘Muller-D’, DX), and its former autosomal homolog (DY) has replaced the ancestral Y chromosome^32^ (**Fig. 1a**). This transition of autosomes to sex chromosomes occurred within the last 15 million years^33^, providing *Dpse* the advantage in dissecting the complex relationships between new genes, sex chromosomes and new cell types which are often difficult to resolve in older sex chromosome systems shaped by long-term evolutionary processes^20,30^. Here by comparative analyses of testis single-cell transcriptomes of *Dpse* vs. *D. melanogaster* (*Dmel*), we seek to address several important questions: 1) How do the new sperm classes of *Dpse* develop in the testes? 2) How is the sex chromosome, the hotspot of new gene birth, dynamically regulated during spermatogenesis? 3) How are the new genes, among other genes, contributing to the evolution of new sperm morphs?

**Fig. 1.**
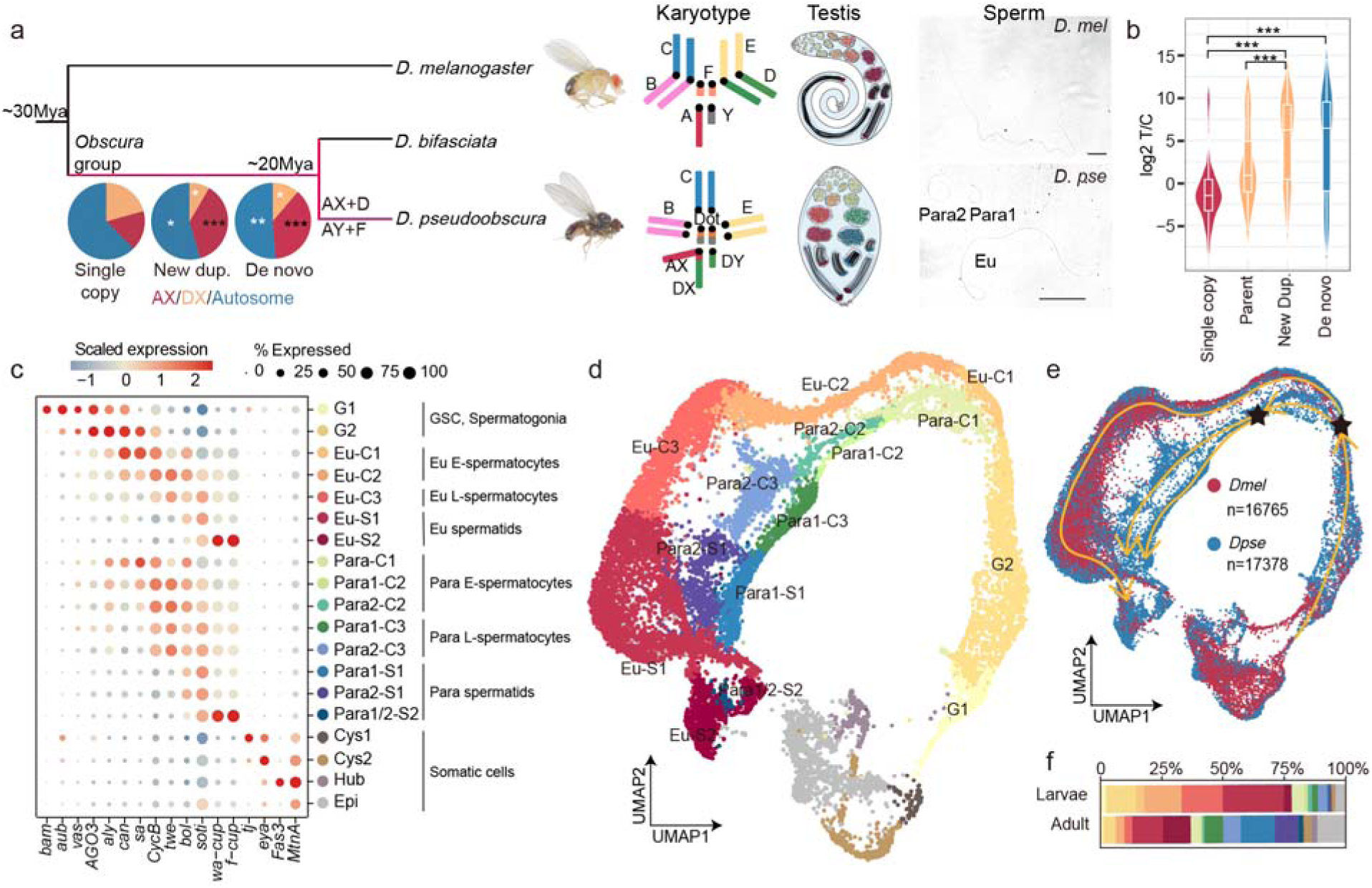
New genes and development of sperm heteromorphism in *D. pseudoobscura*. **a** Phylogeny, karyotype, testis and sperm of *D. melanogaster* and *D. pseudoobscura*. Pie charts show the chromosomal distribution of *Dpse* single-copy genes, lineage specific new duplicated genes and *de novo* genes. The new genes are significantly enriched (black asterisks) on the AX and depleted (white asterisks) on the DX and autosomes (* *P* < 0.05, ** *P* < 0.01, *** *P* < 0.001). The two species’ testes have different shapes, but similar cell type compositions, including germline stem cells and early spermatogonia (G1), late spermatogonia (G2), different stages of spermatocytes (C1-C3), early and late spermatids (S1, S2), hub cells, early and late-stage cyst cells (Cys1, Cys2) and terminal epithelial cells (Epi). See **c** for the color code of each testis cell type. We also show bright-field photos of sperms in seminal vesicles in *Dmel* (top) and *Dpse* (bottom). Scale bar: 100μm. Fly images credit: Nicola White. **b** Violin plots compare the testis specificity (the expression ratio of testis vs. male carcass, T/C) of single copy genes, parental genes, new duplicated genes, and *de novo* genes. *** *P* < 0.001. **c** Expression of marker genes across cell types. Circle size represents the percentage of cells expressing each gene, whereas color indicates the scaled average expression level within each cell type. **d** Uniform Manifold Approximation and Projection (UMAP) of *Dpse* testis scRNA-seq data with cell type annotation. **e** A UMAP integration of *Dpse* and *Dmel* testis cells identifies the *Dpse* specific germ cell lineages. Arrows indicate the trajectory of germ cell development and stars indicate two bifurcation points. **f** Change of cell type composition from larvae to adult testes in *Dpse*.

## Results

### Three classes of *D. pseudoobscura* sperm successively arise during early meiosis

To characterize the contribution of new genes to sperm heteromorphism, we first identified 146 new duplicated genes and 81 *de novo* genes that are present in the *Dpse* genome, but are absent in multiple outgroup *Drosophila* species’ genomes (**Materials and Methods**). We followed the established method of ^34^ to identify *de novo* genes using the syntenic relationships between species. Both new duplicated genes and *de novo* gene that originated either in the ancestor of *obscura* group species or after the species divergence and the sex chromosome turnovers in *Dpse*, are enriched (*P* < 0.001, two-sided Fisher’s exact test) on the *Drosophila* ancestral X chromosome AX, but not on the DX, possibly due to its younger age (**Fig. 1a and Supplementary Fig. 3,** see below). Consistent with previous results^2,7^, new genes show a significantly (*P* < 0.001, two-sided Wilcoxon test) higher level of expression bias toward testis than their parental genes and single copy genes shared with other species (**Fig. 1b**), with their functions in sperm heteromorphism to be characterized below.

We collected the testis scRNA-seq data from the *Dpse* MV2-25 strain 3^rd^ instar larvae and adults with high reproducibility (Person’s correlation *R*=0.97, *P* < 2.2e-16; **Supplementary Fig. 4**). Using a curated list of marker genes^35,36^ (**Fig. 1c**), we annotated a total of 17,378 cells from all collected samples (**Fig. 1d and Supplementary Fig. 5**), with the somatic cells (hub, cyst and epithelial cells) clearly separated from the germ cell lineages in the UMAP projection. *Drosophila* spermatogenesis is a classic model for studying the maintenance and differentiation of adult stem cells^37–40^. The process is initiated by the asymmetric division of each germline stem cell, resulting in one self-renewing stem cell and one goniablast. Goniablast undergoes several rounds (four in *Dmel* or five in *Dpse*) of transit-amplifying divisions^41^ and produces early (G1) and late (G2) spermatogonia cells (**Fig. 1d**). Alignment of *Dpse* testis single-cell transcriptomes against those of *Dmel*^35,36^ (**Fig. 1e**) combined with the RNA velocity analyses (**Supplementary Fig. 6**) identified two critical bifurcation points from which parasperm cell lineages are specified (**Fig. 1e**). The first bifurcation occurs when the spermatogonia cells of *Dpse* develop into two lineages of early spermatocytes (C1), both of which are marked by enriched expression of the well-characterized testis TATA-binding protein associated factors (tTAFs) and testis meiotic arrest complex (tMAC) genes *can*, *sa*, *aly* etc relative to other cell types^42–44^ (**Fig. 1c**). One C1 lineage is clustered with that of *Dmel* based on single-cell transcriptomes, and therefore likely corresponds to the early eusperm spermatocytes (Eu-C1); the other C1 lineage is specific to *Dpse* and likely represents paraspermatocyte progenitors (Para-C1). These progenitor cells undergo the second bifurcation during transition from the early to middle stage of spermatocytes and develop into two lineages of middle-stage parasperm spermatocyte cells (Para1-C2 and Para2-C2). In summary, the *Dpse* eusperm and parasperm differentiate from mitotic cells, whereas the two parasperm classes further differentiate from the early spermatocytes during early male meiosis.

The three lineages of spermatocytes (Eu, Para1, Para2) follow separate developmental trajectories and become respectively the late spermatocytes (-C3), early (-S1) and late (-S2) spermatids (**Fig. 1d, 1e**). Late stage (-S2) spermatids (∼0.8% of all annotated cells, **Fig. 1f**) are difficult to be captured due to the interference from sperm tails during scRNA-seq library preparation^45^, and the lowest number of expressed genes (**Supplementary Fig. 7**). As a result, we were unable to reliably discriminate the late spermatids of Para1 and Para2 based on their transcriptomes. Overall, we estimated a total of 81.5% (14352/17601) of all genes with detectable transcription in at least 3 cells, supporting the widespread genome transcription in testis^2^. Early spermatocytes of all three classes of sperm have the highest portion of transcribed genes (31.8% and 20.9 % on average per cell in larvae and adult) whose products are used for largely, but not entirely transcriptionally silenced (11.1% genes on average) post-meiotic stages (**Supplementary Fig. 7**), consistent with the results reported in *Dmel* and mammals^35,46,47^.

Developmental timing differs between the eusperm and parasperm. In larvae, testes expectedly have more (pre-)meiotic cells, and adults are enriched for post-meiotic cells (**Fig. 1f**). The larvae testes of *Dmel* do not contain spermatids^37,48^, while those of *Dpse* contain small proportions of early and late spermatids, confirmed by both our scRNA-seq (**Fig. 1d and Supplementary Fig. 8a**) and microscopic data (**Supplementary Fig. 8b**). Particularly, eu-spermatocytes (Eu-C1 to Eu-C3) and early eu-spermatids (Eu-S1) comprise the great majority (67.8%) of the germ cells, whereas parasperm germ cells remain arrested at early spermatocyte stages, indicating a substantial delay in parasperm maturation relative to eusperm (**Fig. 1f**). To examine the spatial distribution of different sperm classes during their development, we sequenced one abdomen section of a male adult *Dpse* by the Stereo-seq^49^. By projecting Stereo-seq bins onto our scRNA-seq data (**Materials and methods**), we found an expected spatial distribution of cells during spermatogenesis that spermatogonia are located at one tip of the testis, with parasperm and eusperm cell lineages occupying separate compartments between each other (**Supplementary Fig. 9**). This is consistent with the previous finding that parasperm and eusperm develop in separate cysts within testis^50^.

### X chromosome is silenced and dosage compensated during *D. pseudoobscura* spermatogenesis

The X chromosome is enriched for testis biased new genes (**Fig. 1a, 1b**) that likely function in male meiosis. It is unknown whether these genes are subjected to MSCI in *Dpse*; even in *Dmel*, whether MSCI exists remains highly contentious^36,51–54^. One prediction of MSCI, i.e., an excess of testis-biased genes on the autosomes that escaped from the X through duplication or transposition (termed ‘gene traffic’ in mammals) ^53,55,56^, and the consequential demasculinization pattern of X-linked gene expression^57^, have been widely reported among *Drosophila* species, including the more recently evolved DX of *Dpse*^57,58^. It is also unclear whether canonical dosage compensation mediated by the Male Specific Lethal (MSL) protein complex^59,60^ exists in *Dpse* male germ cells, which was reported to be lacking in those of *Dmel*^61–65^. Elucidating how sex chromosomes are dynamically regulated during spermatogenesis of *Dpse* is critical for understanding the transcriptional program underlying the development of sperm heteromorphism in this species.

MSCI predicts a decreasing transcription of X chromosome vs. autosomes during meiosis. To clarify this, we compared the general transcription level of AX/DX vs. autosomes during spermatogenesis and found that DX- but not AX-linked genes show a decreasing relative transcription through meiosis (*P* < 0.001, two-sided Wilcoxon test) (**Fig. 2a, Supplementary Fig. 10**). The lack of meiotic downregulation on AX can be explained by new genes (**Fig. 1a**): after excluding them, the rest single-copy genes on both X chromosomes consistently showed downregulation through meiosis progression (**Fig. 2b**), providing direct evidence for MSCI in *Dpse*. Such patterns are consistent between larvae and adults, and also between eusperm and parasperm lineages (**Supplementary Fig. 11**). In contrast, new duplicated genes and *de novo* genes on the AX showed a significant (*P* < 0.001, two-sided Wilcoxon test) upregulation during meiosis, which likely has a stronger counteracting effect against MSCI due to enrichment of new genes on the AX but not DX (**Fig. 1a**). The Y-linked genes also display meiotic upregulation (*P* < 0.001, two-sided Wilcoxon test) (**Fig. 2a**), suggesting the absence of MSCI and special regulatory program on the Y during spermatogenesis^66^.

**Fig. 2.**
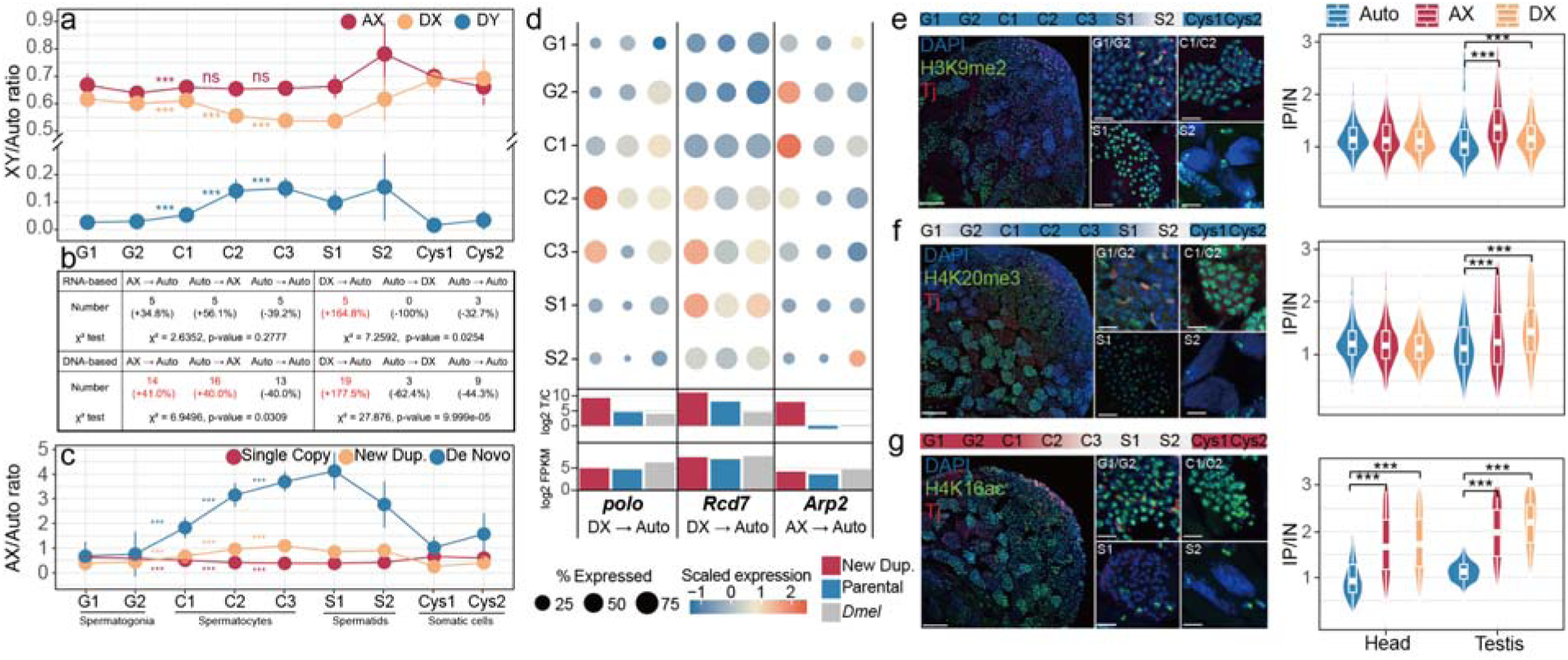
Dynamic regulation of sex chromosome transcription. **a** Relative expression of genes on sex chromosomes versus autosomes. Y-axis is the total scRNA-seq counts from the sex chromosome divided by that from autosomes (Muller B, C and E) normalized by the total number of genes at the single-cell level. Eusperm data is shown here, and parasperm has the same pattern shown in **Supplementary Fig. 10**. **b** Relative expression of AX-linked single copy genes, new duplicated genes or de novo genes compared to autosomes. For (**a**) and (**b**) the expression in adjacent cell types were tested (Wilcoxon two-sided, *** *P* < 0.001). The error bar represents the standard deviation. **c** Table shows the observed number of RNA-based or DNA-based interchromosomal gene movements. Brackets show the excess of gene movement (observed - expected)/expected. Red numbers indicate significant (*P* < 0.05, Chi-square test) enrichment. **d** Single cell and bulk RNA-seq expression profile of new gene cases supporting MSCI. The circle size is scaled to the percentage of cells with expression and the color is the average expression scaled within the same species for each gene group. The spermatocytes (C1-C3) and spermatids (S1, S2) data are from eusperm. **e-g** Testis immunofluorescence (left) and comparison of the gene-level IP/IN (Immunoprecipitation/Input) ratio (right) of H3K9me2 (**e**), H4K20me3 (**f**) and H4K16ac (**g**). For the immunofluorescence panel, the top gradient bar summarizes immunofluorescent results presented below. Panels show the whole-mount apical region of testis (left), spermatogonia (G1/G2), early-to-middle spermatocytes (C1/C2), early spermatids (S1), and late spermatids (S2) (right). Blue: DNA stain DAPI, green: histone marks, red: Early cyst cell marker Traffic jam (tj). Scale bar 50 μm (left) and 10μm (right). Boxes indicate the median and interquartile range within violin plots; \*\*\**P* < 0.001(Wilcoxon two-sided).

MSCI is also supported by an excess (Chi-square test, *P* < 0.05) of DNA- or RNA-based gene movements from the AX and DX onto the autosomes (‘gene traffic’) in the *Dpse* lineage (**Fig. 2c, Materials and Methods**). These relocated new genes onto the autosomes are supposed to escape MSCI and likely have substituted the critical meiotic function of their X-linked parental copies silenced in meiosis. Indeed, relocated genes from the DX are enriched in the Gene Ontology (GO) terms of ‘spermatogenesis’ and ‘centrosome cycle’ (**Supplementary Fig. 12**) and also are more likely than those between autosomes to have an increased spermatocyte expression (one-sided Fisher’s exact test, *P* = 0.0105) compared to their parental genes (**Supplementary Fig. 13**). For example, in *Dmel*, *Rcd7* and *polo* are located on chr3L or the D element and were reported to be respectively required for male fertility and meiotic spindle assembly^67–70^. Another case *Arp2* on the AX, one of the actin related protein (Arp) superfamily genes, produced a RNA-based duplicated copy *Arp2D* onto the autosome in *Dpse*. *Arp2D* was recently reported to show sequence signatures of positive selection, and function in actin cones crucial for the sperm individualization^71^. The *Dpse* autosomal new gene copies, but not the AX/DX orthologous genes of *Rcd7*, *polo* and *Arp2*, show a similar transcription pattern with the *Dmel* genes enriched among spermatocytes or spermatids (**Fig. 2d**). Conversely, genes translocated or duplicated from autosomes onto the AX more frequently (*P* = 0.0138, one-sided Fisher’s exact test, **Supplementary Fig. 13**) have reduced levels of germ cell transcription, bulk testis transcription, or testis specificity compared to their parental copies, like *Naa20A* and *CG31465* (**Supplementary Fig. 14**), supporting MSCI’s contribution to demasculinization of X-linked gene expression. Both chromosome-wide and individual spermatogenesis genes’ transcription patterns support that MSCI is operating on both the AX and DX in *Dpse*.

To reveal the regulatory mechanism of MSCI, and clarify whether there is canonical dosage compensation in male germ cells of *Dpse*, we next performed chromatin immunoprecipitation sequencing (ChIP-seq) and/or testis immunofluorescence (IF) analyses, using antibodies targeting the constitutive heterochromatin marks histone H3 lysine 9 di- and tri-methylation (H3K9me2/3) and H4K20me3, as well as one of the dosage compensation complex proteins MSL-2^72,73^ and its induced active histone mark H4K16ac^74,75^. We found that both AX and DX genes show a significantly (*P* < 0.001, two-sided Wilcoxon test) higher normalized ChIP binding levels of H3K9me2 and H4K20me3 than autosomal genes, specifically in testis but not in male head (**Fig. 2e and 2f**). This is different from the reported results in *Dmel*^63^, whose spermatocytes do not exhibit an enrichment of H3K9me2 on the X chromosome. All three repressive marks show robust IF staining in somatic cells and most germ cells, but start to decay in signals from early spermatids and become completely undetectable in late spermatids during the replacement of histones by protamine (**Fig. 2e, 2f and Supplementary Fig. 15**). Such testis specific enrichment of repressive histone marks on the X chromosomes provides epigenomic evidence supporting that both AX and DX undergo MSCI during the male meiosis in *Dpse*.

The pattern of canonical dosage compensation within the germline also seems to be different between *Dpse* and *Dmel*. Previous research using IF^61^, RNA *in situ* hybridization^62^, and ChIP-seq^63^ failed to detect MSL-2 and H4K16ac stainings, or an enrichment of H4K16ac binding on the X chromosome in germ cells in *Dmel*. scRNA-seq also reported a very low level of transcription of all MSL complex genes in male germ cells of *Dmel*^62^. By contrast, we found that all the MSL complex genes^76,77^ are transcriptionally active in early spermatogonia of *Dpse*, with expression levels above the median of all genes (CPM > 0.102) (**Supplementary Fig. 16**). Most of them remain transcriptionally active until the early-stage spermatocytes except for *msl-2* and *roX1*, which have been silenced in even earlier stages (**Supplementary Fig. 16**). This is similar to the recent result reported in the sister species of *Dpse D. miranda*^52^. We did not find significant differences between eupserm and parasperm cells of the same stage among these MSL complex genes, except for *roX2* (fold change -0.63, *P* < 0.001, two-sided Wilcoxon test) (**Supplementary Fig. 16**). Importantly, our IF staining with MSL-2 and H4K16ac antibodies confirmed that both have binding signals in spermatogonia and early spermatocytes of *Dpse* (**Fig. 2g and Supplementary Fig. 17**). And H4K16ac ChIP-seq patterns indicated a specific enrichment (*P* < 0.001, one-sided Wilcoxon test) on the AX and DX chromosomes relative to autosomes in both testis and head (**Fig. 2g**). In summary, we provided evidence supporting that, different from *Dmel*, both MSCI and canonical dosage compensation are present in the *Dpse* male germ cells.

### Differentiation between sperm classes is regulated by meiotic arrest genes

To dissect the hierarchical gene regulatory networks underlying development of the *Dpse* sperm heteromorphism, we characterized the dynamical network biomarkers (DNB) which are supposed to become more correlated with each other with increased fluctuation of expression, when the biological system is approaching a transitional state. This method has been widely used to identify the pre-disease state before irreversible deterioration, or the tipping point of cell fate transition^78–81^. Taking both the single-cell level expression variation and correlation with other genes into account, a DNB index can be calculated for each gene within each cell type (**Materials and Methods**)^79^. The top 500 genes ranked by their DNB indices were defined as ‘DNB genes’ (DNBGs’) and were then used to produce a DNB index landscape through the *Dpse* spermatogenesis, where a higher average value of DNB indices predicts certain cell type being close to the transition point. The DNB landscape in both *Dpse* larvae and adult testes confirmed the late spermatogonia and early para-spermatocyte as the two critical cell bifurcation time points (**Fig. 1e**), each of which shows a significantly higher (*P* < 2.2e-16, one-sided Wilcoxon test) DNB index of the DNBGs than that of adjacent stages (**Fig. 3a and Supplementary Fig. 18**). By contrast, the DNB landscape of *Dmel* without sperm heteromorphism does not show a similar increase of DNB index at the two corresponding stages during spermatogenesis. Likely as a result of MSCI, DNBGs and their putative downstream differentially expressed genes between eusperm and parasperm (EP-DEG, see below) are depleted (*P* < 0.01, one-sided Fisher’s exact test) on the DX and/or AX **(Supplementary Fig. 19)**. Interestingly, the DX-linked DNBGs of late spermatogonia show a significantly (*P* = 7.065e-05, two-sided Wilcoxon test) higher binding level of active mark H3K4me3, but lower (*P* < 0.01, two-sided Wilcoxon test) binding levels of repressive marks H3K9me2 and H3K20me3 than other genes of the same chromosome, and a similar (*P* > 0.05, Wilcoxon test) binding level of repressive marks with autosomes (**Supplementary Fig. 20**). This suggests that some X-linked DNBGs might have escaped the MSCI.

**Fig. 3.**
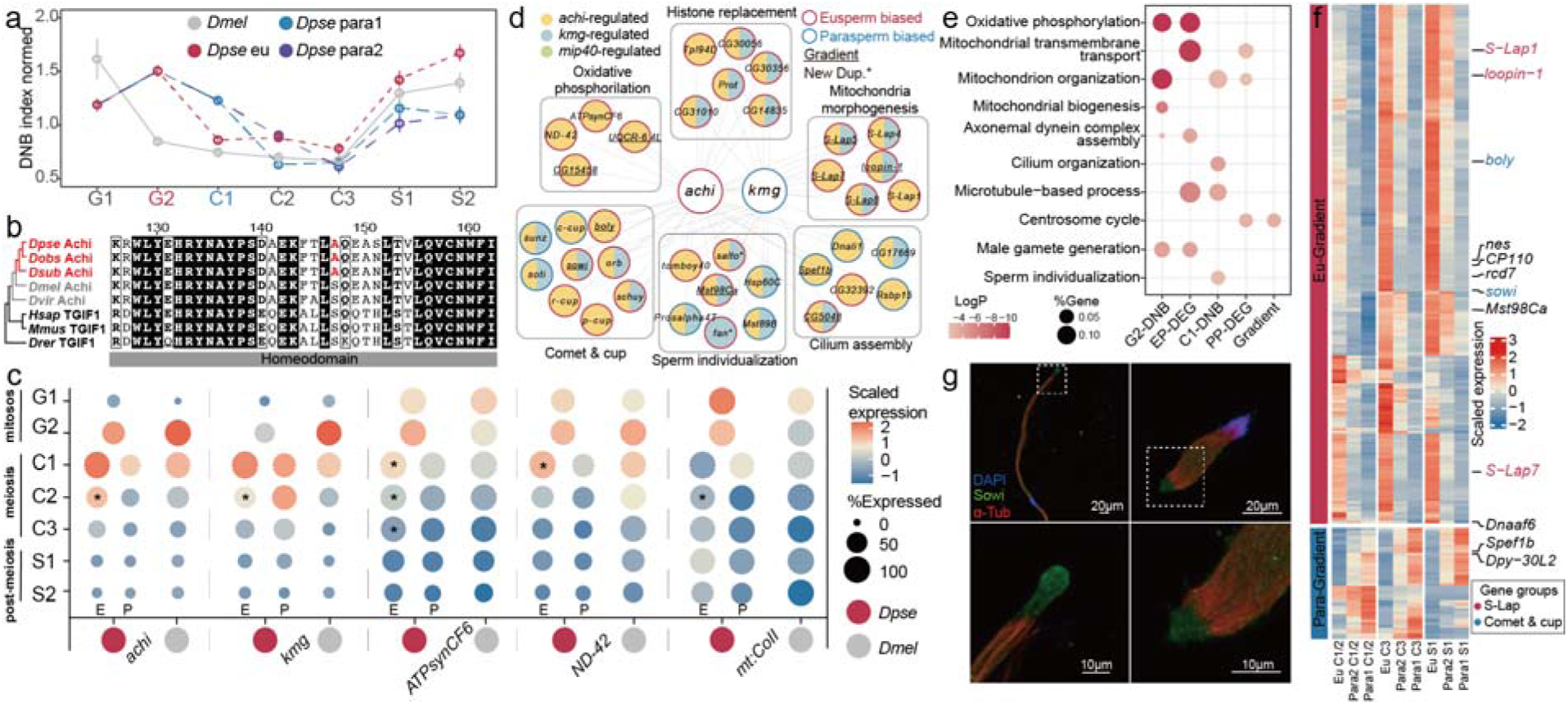
Transcriptional regulation underlying differentiation of eusperm and parasperm in *Drosophila.* **a** Normalized DNB index at each cell type. Dot shows the average DNB index of the top 500 genes normalized by the average DNB index of all genes. The error bar represents the standard deviation relative to the mean value. Adult data is shown here, and larval data has the same pattern shown in **Supplementary Fig. 18. b** Alignment of part of Achi homeodomain region from *Drosophila obscura* group species (red), other *Drosophila* species (grey) together with TGIF1 homologs from vertebrates (black). The red amino acid indicates *obscura*-group–specific substitution in the Achi proteins. The full homeodomain sequences are shown in the **Supplementary Fig. 20. c** Bubble plot showing differential expression of meiotic arrest genes (*achi*, and *kmg* or genes in mitochondrial complexes (*ATPsynCF6*, *ND-42* and *mt:COII*). E, eusperm; P, parasperm. The asterisk (*) indicates the expression level in this cell morph is higher than that in the other cell morph at the same stage. **d** Putative regulatory network of DEGs associated with *achi* and *kmg*, inferred from loss-of-function transcriptome analyses or ChIP experiments^77,79^ of orthologous genes in *Dmel*. **e** GO enrichment of DEGs, DNBGs and gradient genes. **f** Expression profile of gradient genes. Genes with spermatogenesis functions were labeled. *S-Lap* genes are shown in red while comet & cup genes are shown in blue. The expression value is scaled by z-score. **g** Immunofluorescence of *sowi* in eusperm (left) and parasperm (right). Blue: DNA stain DAPI, green: *sowi*, red: α-Tubulin.

Four *Dpse* orthologs of the 12 reported ‘meiotic arrest’ genes in *Dmel* were identified as DNBGs in late spermatogonia and early para-spermatocytes. *Dmel* mutants of these master regulators of spermatogenesis show meiotic arrest and significant changes of expression of over 1000 genes in testis^38,82^. The four DNBGs include two testis-specific meiotic arrest complex (tMAC) genes *achi*, *wuc*^44,83^, two testis TATA-binding protein associated factors (tTAFs) *nht* and *rye*^84^. In addition, the tMAC regulator *kumgang* (*kmg*) ^85^ was also identified as a DNBG. We found that the amino acid sequence of the homeodomain region of *achi*^86^ is extremely conserved across animals, except for a non-synonymous mutation shared by all *Drosophila obscura* group species (**Fig. 3b and Supplementary Fig. 21**). This might alter *achi’*s DNA-binding capacity specifically in these species all with sperm heteromorphism. In addition, DNBGs *achi* and *kmg* evolved biased (log2(fold-change)>0.5) expressions toward eu-spermatocytes or para-spermatocytes (**Fig. 3c**). Such biased expression likely constitutes one of the key upstream signals orchestrating many other downstream DEGs between eusperm and parasperm (EP-DEG, 1274 or 8.9% of transcribed testis genes), and between the two types of parasperm (PP-DEG, 233 or 1.6% of transcribed testis genes). Indeed, 50.8% of the EP-DEGs, and 43.3% of the PP-DEGs in *Dpse* are orthologs of the known downstream genes regulated by *achi* and *kmg* in *Dmel* inferred by their loss-of-function or ChIP experiments^85,87^ (**Fig. 3d**). In addition, DNBGs in late spermatogonia, and EP-DEGs have a significantly (*P* < 0.05, two-sided Wilcoxon test) faster evolution rate (measured by the ratio of nonsynonymous vs. synonymous substitution rate, dN/dS) in coding regions than their *Dmel* orthologs (**Supplementary Fig. 22**). These results indicate changes of both protein sequences and gene expression are involved in evolution of the regulatory network consisting of DNBGs and DEGs that is responsible for developing sperm heteromorphism.

### Eusperm and parasperm differ in energy investment and sperm morphogenesis genes

Both DNBGs and eusperm biased, but not parasperm biased genes are enriched (*P* < 0.05, hypergeometric test) in GO terms of energy metabolism (‘ATP metabolic process’, ‘mitochondrial transmembrane transport’ etc., **Fig. 3e and Supplementary Fig. 23**). This supports the hypothesis that parasperm are ‘cheaper’ in terms of energy cost to be produced^24,26,88^. They possibly protect the eusperm through their higher sensitivity to and thus higher efficacy in scavenging reactive oxygen species in the male or female reproductive tract^89,90^. This is implicated by the pattern that parasperm biased genes are enriched in GO terms of ‘cellular response to oxidative stress’ (**Supplementary Fig. 23**). Particularly, the eusperm biased genes are overrepresented (*P* < 0.05, one-sided Fisher’s exact test) among the five mitochondrial respiratory chain complexes (with a total of 113 orthologous genes) that generate the majority of cellular energy through oxidative phosphorylation. 34 or 89.5% of the DEGs encoding mitochondrial complexes have a higher (log2FC > 0.5) expression in eusperm than in parasperm in at least one cell stage. They include both the genes on the mitochondira *mt:COII/COIII*, and the *achi*-regulated nuclear genes *ND-42* and *ATPsynCF6* (**Fig. 3c and 3d**) whose male mutants are infertile^91–93^. The reduced expression of these genes could be one contributing factor to the sterility of parasperm, as some of their mutants in *Dmel* show defects in spermatid individualization (e.g., *COII*) ^91^ or elongation (e.g., *ND-42*^92^).

In fact, DNBGs in late spermatogonia are significantly (Bonferroni-corrected *P* < 0.05, Fisher’s exact test) enriched for *Dmel* mutant phenotypes in spermatid elongation and individualization that involves morphogenesis of nuclei in the spermatid head, and that of mitochondria in the spermatid tail^94^. It is hypothesized that the wider head of *Dpse* parasperm than eusperm could prevent them from passing the micropyle of egg for fertilization^95^, consistent with the reported different nuclear compaction level between eusperm and parasperm^22,96^. These head morphological differences might be associated with the DEGs encoding sperm nuclear basic proteins (SNBPs) that replace the histones for packaging the genome in the sperm nuclei. 6 of the total 10 SNBP genes annotated in *Dpse*^97^, including *Protamine* (*Prot*), show eusperm biased expression, and all these genes are regulated by *achi*, *kmg* or *mip40* (**Fig. 3d**).

Besides head morphology, differences in the tail length and potentially motility between sperm classes can be explained by 229 EP-DEGs (229 or 17.9% of EP-DEGs) that exhibit a gradient of transcription levels corresponding to that of tail lengths among the three sperm classes (**Fig. 3f**). These gradient genes include 186 genes that have the highest expression in eusperm with the longest tail, and the lowest expression in Para1 with the shortest tail, and 43 genes showing a reversed gradient. These two combinations of gradients are overwhelmingly (Chi-square test, *P* < 0.001) abundant among all permutations of the three classes of sperm. Particularly, 10 of the identified 17 *Dpse* orthologs of *Dmel* ‘comet’ and ‘cup’ genes^98^ are gradient genes or EP-DEGs and 9 of them are inferred to be regulated by *achi*, *kmg* or *mip40* in *Dmel* (**Fig. 3d and 3f**). The ‘comet’ and ‘cup’ genes could be linked to the tail elongation given their localized expression to the distal ends of maturing spermatids^98^. We confirmed the eusperm-biased expression pattern for one of the ‘comet’ genes *sowi* in *Dpse* through immunofluorescence (**Fig. 3g**). The eusperm biased genes or gradient genes also include 6 of the 10 annotated Sperm-Leucylaminopeptidases or *S-Lap* genes whose protein products accumulate within the elongated mitochondrial derivatives and play a structural role in sperm tail elasticity and undulation^99,100^ (**Fig. 3d, 3f**). In conclusion, we have reconstructed a complex gene regulatory network centered on a few meiotic arrest genes, which likely initiates the differential expression of genes involved in energy metabolism, and head/tail morphogenesis between eusperm and parasperm (**Fig. 3d**).

### New genes play a prominent role in the emergence of new sperm classes

Both the *de novo* genes and new duplicated genes specific to the *Dpse* lineage are enriched (*P* < 0.001, one-sided Fisher’s exact test) for the upstream DNBGs and the downstream DEGs, including the identified gradient genes, compared to all the transcribed testis scRNA-seq genes as a background (**Fig. 4a**). In addition, these new genes also show a significantly (*P* < 0.05, Wilcoxon test) higher extent of differential expression between eusperm and parasperm than the other single-copy genes (**Fig. 4b**). These results provide evidence that the new genes play a more prominent role in the emergence of new sperm classes throughout their entire regulatory program. Why are the new genes more likely than other genes to contribute to evolution of new sperm classes? Such propensity probably stems from their distinctive transcriptional patterns compared to their parental copies or other single-copy genes.

**Fig. 4.**
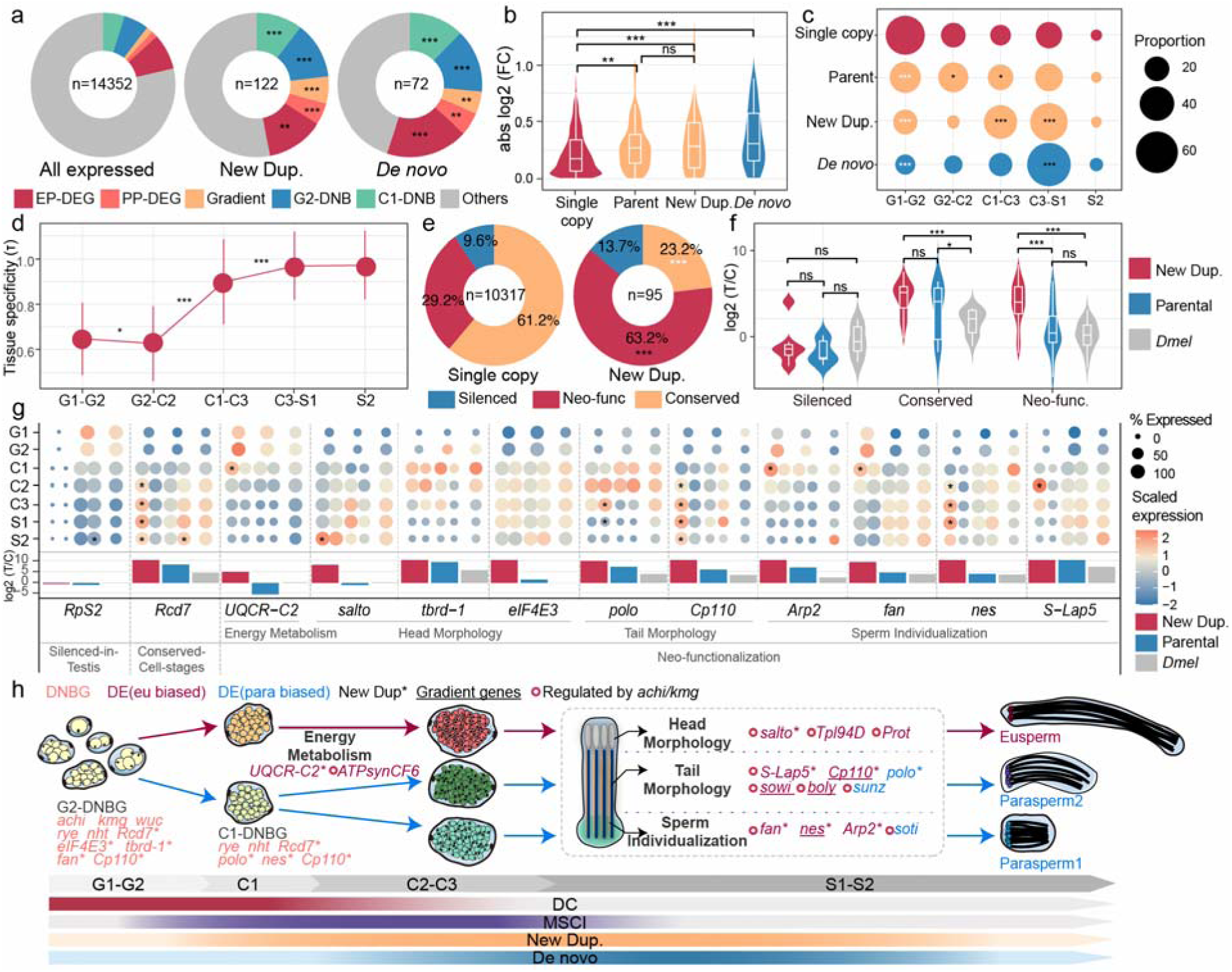
function of DE and DNB genes and the molecular mechanism of sperm heteromorphism. **a** Donut plots show the proportion of genes assigned to each expression category (EP-DEG, PP-DEG, Gradient, G2-DNB, C1-DNB, and Others) within all expressed genes, newly duplicated genes, and de novo genes. **b** New genes show a significantly (*P* < 0.05, Wilcoxon test) higher extent of expression difference between eusperm and parasperm compared to single-copy genes, with the maximum absolute fold change across cell stages used for each gene. **c** Distribution of the predominantly expressed testis cell types of each category of genes. The predominantly expressed cell types were annotated through hierarchical clustering (**Supplementary Fig. 24**). **d** Tissue specificity index (τ) of sets of genes predominantly expressed within each testis cell type. **e** Donut plots show the proportion of genes classified as silenced-in-testis, conserved-cell-stages, or neo-functionalized among single-copy genes and new duplicated genes. **f** Testis specificity (T/C) for new duplicated genes, parental genes and *Dmel* orthologs across three functional categories. **g** Expression of new duplicated genes, parental genes and *Dmel* orthologs at single-cell and bulk RNA-seq level. For *Dpse* genes, the left and right column of bubbles shows eusperm and parasperm respectively. The asterisk indicates the expression level in this cell morph is higher than that in the other cell morph at the same stage. **h** Inferred hierarchical gene regulatory networks highlighting the roles of new genes in *Dpse* sperm heteromorphism. In (**a**), (**c**) and (**e**) asterisks indicate statistically significant enrichment (black) or depletion (white) compared to all genes or single copy genes (one-sided Fisher’s exact test). In (**b**), (**d**) and (**f**) asterisks indicate differences between groups, (two-sided Wilcoxon tests). * *P* < 0.05, ** *P* < 0.01, *** *P* < 0.001.

By annotating each transcribed testis gene with their dominantly expressed male germ cell types through hierarchical clustering (**Supplementary Fig. 24**), we were able to compare the cell-level expression patterns of new genes through spermatogenesis and against their parental genes. Compared to the parental or single-copy genes, the general expression of new duplicated genes is more concentrated (*P* < 0.001, one-sided Fisher’s exact test) in spermatocytes and early-stage spermatids, while that of *de novo* genes is shifted toward late spermatocytes and spermatids (**Fig. 4c**). This expression propensity toward late or after meiosis explains why new genes on the X chromosome manage to escape MSCI (**Fig. 2b**). Genes (not only new genes) with concentrated expression in late-stage spermatocytes and early-stage spermatids have a significantly (*P*<0.001, one-sided Wilcoxon test) higher extent of tissue-specificity (τ) in expression, compared to the genes with concentrated expression in cells of earlier spermatogenesis stages (**Fig. 4d**). This supports the hypothesis that the permissive transcription chromatin state during and after male meiosis facilitates the birth of new genes. Because genes expressed at these stages tend to have a very confined expression to testis, potentially minimizing deleterious pleiotropic effects on other tissues and increasing their likelihood of retention. The expression propensity of *Dpse* new genes is consistent with previous results reported in *Dmel* and mammals^34,35,101,102^, and overlaps with the stages during which new sperm classes develop. It therefore predisposes new genes to be more often than other genes involved in the emergence of sperm classes.

Another key factor underlying the prominent role of new genes in new sperm classes origination is the frequent functional innovation of newly duplicated genes^103^. By comparing the testis cell-stage expression of new duplicated genes vs. those of parental genes and *Dmel* orthologs, we classified 95 *Dpse* lineage specific gene duplications into three functional categories (**Fig. 4e**): i) conserved cell stages, in which both copies retain similar expression patterns; ii) neo-functionalization, in which one copy acquires a novel cell-stage expression pattern or acquired testis specificity; and iii) silenced-in-testis, in which one copy shows no detectable transcription across all testis cell stages (**Materials and methods**). Besides 13.7% of the new gene duplicates have become silenced probably of their dosage constraints on housekeeping functions (e.g., the ribosomal proteins *RpS2/3*, **Fig. 4g**). 63.2% of them showed signatures of neo-functionalization that are expressed in new testis cell stages, or evolved testis-specific expression pattern (**Fig. 4e**). Accordingly, new duplicated genes display a generally higher testis specificity than their parental genes (**Fig. 4f**), providing direct evidence that new genes are more likely to undergo neo-functionalization than other testis genes.

### New genes have putative functions involved in morphogenesis of sperm head and tail

New duplicated genes can rewire the ancestral spermatogenesis network to accommodate the development and functional differentiation of new sperm classes, by expressing in the same or in the new testis cell types compared to their parental copies. For example, *Rcd7* is required for centrosome separation in spermatogenesis of *Dmel*^68,104^ and presumably required for development of all sperm classes in *Dpse*. Although the two copies of *Dpse Rcd7* have similar expression across cell stages, the *Dpse*-specific duplication was identified as a DNBG and a eusperm biased gene, while the parental copy was not a DNBG and showed parasperm biased expression (**Fig. 4g**). This suggested that gene duplicate pairs in *Dpse* could partition their ancestral functions between the different sperm classes. Some other duplicate genes evolved novel testis-specific expression, as illustrated by UQCR-C2 with a role in mitochondrion electron transport^105^. Compared to its broadly expressed parental copy, its duplicated copy in *Dpse* showed biased expression in testis and also in eusperm (**Fig. 4g)**, supporting again more energy investment in eusperm development.

For the new gene duplicates that evolved a new expressed testis cell stage, or neo-functionalization, they are enriched (*P* < 0.01, Hypergeometric test) in GOs of spermatogenesis, cell cycle and centrosome cycle **(Supplementary Fig. 25)**. These genes include the aforementioned *Arp2* (**Fig. 2D**), as well as *fan*^106^ and *nes*^107^, all of which are known in *Dmel* to function in sperm individualization. The *Dpse*-specific duplicates of all these genes evolved a eusperm biased expression (**Fig. 4G**). Parental copies of other new duplicated genes have more direct implications for the morphological and functional divergence between the sperm classes. Mutants of *salto*^108^, *tbrd-1*^109,110^, and *eIF4E3*^111^ in *Dmel* were shown to have changed their nuclei into coiled, small and round, or other defected shapes relative to the wildtype. And mutants of *polo*^112^, and *Cp110*^113,114^ exhibit extra or shorter axonemes or elongated centrioles. In particular, *Dpse*-specific duplicate of *S-lap5*, whose parental gene encodes paracrystalline material accumulated in mitochondrial derivatives of sperm tail^99^, evolved a eusperm biased expression. In summary, new duplicated genes emerged among both upstream DNBGs and downstream DEGs and likely have impacted the head and tail morphological difference between sperm classes of *Dpse* (**Fig. 4h**). In future, the functions of *de novo* genes that constitute a disproportionately high number among DNBGs and DEGs as well as new duplicated genes (**Fig. 4a**) will be of particular interest to be elucidated.

## Discussion

Spermatogenesis is an ancient developmental process with a deeply conserved genetic toolkit shared across metazoans^4^. However, during the late stages, different species produce spermatids that vary tremendously in number and morphology^18,115^. At the same time, the past decades of new gene studies have well documented that this stage comprises a hotspot for new gene expression and birth, due to its distinctive transcriptional program and intensive sexual selection^116^. Our work here provides a pivotal link between the evolutionary innovations at the cell and genetic levels during spermatogenesis in *Dpse*, and have broad implications for evolution of cellular altruism.

We found *Dpse* spermatogenesis has a distinctive transcriptional regulation and developmental trajectories compared to that of *Dmel*. We provided evidence from chromosomal histone modification patterns and analyses of X-linked ‘traffic genes’ that *Dpse* undergoes MSCI and dosage compensation during mitosis and early meiosis (**Fig. 4h**), which are absent or under debate in male germ cells of *Dmel*^36,51–54,61–65^. Many X-linked new genes can escape MSCI, probably because it is a general propensity of new genes, regardless of their chromosomal locations (**Fig. 4c** and **Supplementary Fig. 26**), to be transcribed toward late and after meiosis^34,35,101,102^ (**Fig 2b**). Interestingly, Y-linked genes have a similar transcription trend (**Fig 2a and Supplementary Fig. 27**). This shared propensity could be associated with the fact that substantial numbers of both Y-linked genes and new genes reside in heterochromatin (**Supplementary Fig. 28**) that only become lost during the histone-to-protamine transition in spermatids. It remains unclear how Y-linked genes and new genes, as well as the ‘comet and cup’ genes located to the distal ends of elongating spermatids^117^, are transcribed in the predominantly transcriptionally silent spermatids. A recent study suggested that many new genes in *Dmel* are *de novo* activated in mid-to-late elongating spermatids, whereas others are from residual transcription in spermatocytes^47^. Such late-stage transcriptional activation may more frequently expose new genes to sperm competition in *Dpse*, thereby contributing to the emergence of new sperm classes and the diversification of morphology and functions relative to eusperm.

The over 200 identified genes enriched for new genes that form a gradient of transcription levels between sperm classes (**Fig. 3h**) constitute a key dataset for future functional validation of the genetic factors impacting the lengths and mobility of sperm tails. Several candidate upstream regulators of these gradient genes include *Rcd7*, *nes* and *Cp110*^68,107,118^, whose *Dmel* orthologs have known functions in spermatogenesis and their new gene copies, but not parental copies in *Dpse* are identified as both a DNBG and a gradient gene (**Fig. 4h**). In addition, other identified DEGs associated with the sperm head formation (e.g., *Prot* and *salto*) could account for the distinct head morphology^22,96^. *Salto* is a recently duplicated gene that becomes enriched at the acrosome in fully elongated spermatids, suggesting a role in acrosome specialization underlying the lack of fertilization capacity of parasperm. Overall, as nearly 50% of these DEGs between sperm classes have *Dmel* orthologs regulated by a few meiotic arrest genes (e.g., *achi* etc.), the hierarchical regulatory network of sperm heteromorphism in *Dpse* is likely co-opted from an ancestral network centered on the meiotic arrest genes that is shared with *Dmel*^44,84^(**Fig. 3e**).

From both mitochondrial and nuclear genes, we found consistent evidence supporting that parasperm development involves less energy investment with a lower transcription level from genes of energy metabolism. Thus, parasperm likely act as a ‘cheap shield’ protecting the eusperm during fertilization. Suggested by previous close inspection of the relative percentage between eusperm vs. parasperm across different time points after mating^119^, this protection occurs only within the female storage organ when female cryptic selection may also occur. Here we have identified differentially expressed genes between sperm classes, but how these genes’ protein products differentially interact with other male (e.g., accessory gland) and female (e.g., spermatheca) proteins should be targeted in future studies to understand the mechanisms of how parasperm sacrifices their primary function to protect the eusperm.

## Materials and Methods

### Genome annotation and sperm inspection

The improved *Dpse* genome assembly and annotation was generated and described in ^120^. by combining a female reference assembly (UCI_Dpse_MV25, https://www.ncbi.nlm.nih.gov/datasets/genome/GCF_009870125.1/) with Y-linked scaffolds from a male assembly (UCBerk_Dpse_1.0, https://www.ncbi.nlm.nih.gov/datasets/genome/GCA_004329205.1/). The Y chromosome was connected into a chromosome using Hi-C data, and Y-linked gene annotation was performed with MAKER v2.31.10^121^. We totally identified 12022 pairs of *Dmel*-*Dpse* 1:1 orthologous gene using reciprocal best hits from pairwise BLASTP (v2.2.26) searches with an E-value cutoff 1e-7. The orthologous genes were used for cross-species integration of scRNA-seq data.

We used OrthoFinder v2.5.2^122^ to construct orthogroups with all protein sequences from 5 *Drosophila* species: *D. melanogaster* (FlyBase r6.15), *D. virilis* (NCBI GCF_003285735.1), *D. willistoni* (NCBI GCF_000005925.1), *D.subobscura* (NCBI GCF_008121235.1) and *D. pseudoobscura*. Genes present as a single copy in *Dpse*, *Dsub* and *Dmel* within the same orthogroup are defined as single copy genes. *De novo* genes were identified following the method from^34^ based on the conserved synteny. We built Cactus^123^ whole-genome alignments from 11 *Drosophila* species (*D. ananassae*, *D. erecta*, *D. grimshawi*, *D. melanogaster*, *D. mojavensis*, *D. pseudoobscura*, *D. simulans*, *D. suzukii*, *D. virilis*, *D. willistoni*, *D. yakuba*), and applied halLiftover^124^ to map syntenic regions of *Dpse* genes to other species. *Dpse* genes whose syntenic loci in all other species were (i) unannotated, and (ii) had <30% sequence identity and <30% alignment coverage when searched by BLAST (E-value cutoff 1e-6) against protein or CDS sequences of other species and against UniProt, were considered candidate *de novo* genes. We further applied Genewise^125^, and Spaln^126^ to predict protein coding potential in syntenic regions, and excluded candidates whose synteny regions could be putative protein-coding genes. In total, 81 *Dpse* de novo genes were identified.

To inspect the sperm, *Dpse* stocks (MV2-25, NDSSC stock # 14011-0121.94) were raised on standard Bloomington medium at 18L with a 12-hour light/dark cycle. Seminal vesicles from 5- to 7-day-old adults were dissected in cold phosphate buffer saline (PBS) and then ruptured and dragged several times on a glass slide to release sperm. Sperm were visualized on Nikon QS-Qi2 microscope at 200x magnification. For each pair of seminal vesicles, we took 20 images with NIS-Elements BR 4.60.00 and measured the length of all observed sperms by ImageJ (https://imagej.nih.gov/ij/) (43-154 sperms per male, average 106 sperm, N=10). 10 individual adults were randomly selected for calculation.

To examine sperm tails by transmission electron microscopy, testes from 5- to 7-day-old adults were dissected and fixed overnight at 4 CC in 2.5% glutaraldehyde in 0.1 M phosphate buffer (pH7.0). Samples were washed three times in the phosphate buffer for 15 min each, post-fixed in 1% OsO4 in the phosphate buffer for 1.5 h, and washed again three times for 15 min each. Samples were dehydrated through a graded series of ethanol (30%, 50%, 70%, 80%) followed by a graded series of acetone (90%, 95%) for 15 min at each step, then dehydrated twice in absolute acetone for 20 min each. Samples were infiltrated in a 1:1 mixture of absolute acetone and Spurr resin for 1h at room temperature, transferred to a 1:3 mixture of absolute acetone and the resin for 3h and finally placed in pure Spurr resin overnight. Samples were then placed in Eppendorf tubes containing Spurr resin and polymerized at 70L for more than 9h. Ultrathin sections were prepared using a LEICA EM UC7 ultramicrotome, stained with lead citrate solution and uranyl acetate solution for 10 min each respectively and examined in Hitachi Model H-7650 TEM.

### Transcriptome data generation and processing

For *Dpse* samples, each adult sample used 20 pairs of testes from 5- to 7-day-old males and each larval sample used 20 pairs of testes from third instar larvae. We performed 2 replicates for each stage. Testis were dissected in cold PBS and immediately digested in 200ul lysis buffer (HBSS+5%FBS+2mg/ml collagenase IV) for 10 min at room temperature. After digestion, cells were filtered through a 30 μm filter and centrifuged 6 min at 600x g. Cells were washed twice in 200 ul cell suspension buffer (HBSS+5%FBS+0.04%BSA), centrifuged 6 min at 500x g and then resuspended with 100 ul suspension buffer. A 5 μl aliquot of cell suspension was mixed with trypan blue dye to examine cell viability and concentration. We got adult testis cell suspension with viability of 70%-76% and L3 larval testis cell suspension with 81%-89% viability. Cell concentration was adjusted to 1000-1200 cells/μl and samples were immediately proceeded with the 10x Genomics Single Cell 3’ v2 workflow according to the manufacturer’s protocol. Libraries were sequenced onIllumina NovaSeq 6000. Each sample was sequenced for 120Gb.

To generate single-cell expression matrices, we converted Illumina Binary Base Call (BCL) files into fastq using Cellranger 4.0.0 mkfastq (https://support.10xgenomics.com/single-cell-gene-expression/software/pipelines/latest/what-is-cell-ranger) and performed alignment, filtering, barcode counting, and UMI counting using Cellranger count with default parameters. The *Dmel* single-cell datasets were obtained from previously published studies^35,36^. For the reference genome and annotation, we used FlyBase r6.15 (http://ftp.flybase.net/genomes/dmel/dmel_r6.15_FB2017_02/) for *Dmel* and improved assembly and annotation for *Dpse*. Potential doublets were detected using DoubletFinder^127^ and removed. For DoubletFinder the number of statistically-significant principal components (PCs) were set as 30, and the number of artificial doublets (pN) was set as 0.2. The expected doublet rates were calculated based on cell recovery for each sample. Then we used R package Seurat 4.1.1^128^ to perform cell filtering, clustering, and integration. First, genes expressed in at least 3 cells and cells with at least 3000 UMIs were kept for *Dpse* samples. For *Dmel* larval and adult samples, the filtering was performed as described in ^35,36^. We normalized data with command NormalizeData with default parameters. To integrate *Dmel* and *Dpse* cells, the top 2,000 most variable species-orthologous genes were used for canonical correlation analysis (CCA) integration^129^ and 3 neighbors (k) were used when picking anchors. Cells were visualized with Uniform Manifold Approximation and Projection (UMAP) using 30 Principal Components (PCs).

To generate testis Stereo-seq data, abdomen segments were dissected from 5- to 7-day-old *Dpse* adult flies in cold PBS. Dissected samples were rinsed in PBS and dried with wipers. Samples were embedded in Tissue-Tek OCT and immediately transferred to an -80L freezer for storage. Cryosections were cut at a thickness of 10 mm in a Leika CM1950 cryostat at -20L. Stereo-seq library preparation was performed following the procedure described in^49^ and sequenced was performed using a MGI DNBSEQ-Tx sequencer.

We processed the raw data using standard workflow SAW (https://github.com/STOmics/SAW) to generate the CID-containing expression profile matrices^49^. The matrices were binned into non-overlapping 50x50 DNB bins. We analyzed Sterero-seq data with R package Seurat 4.1.1^128^. Data were normalized with SCtransform, followed by FindNeighbors and FindClusters with 30 PCs and a resolution of 1. To annotate the major tissue types, clusters enriched for tissue-specific markers were identified: testis (e.g., *Mst98Ca*^130^ and *eIF4E3*^111^), epithelium (*MtnA*^131^ and *Obp44a*^132^), fat body (*CG30008*^133^ and *Npc2g*^134^), gut (*Ag5r2*^135^ and *Tsp42Ec*^136^), malpighian tubule (*CG10513*^137^ and *CG3270*^138^), and muscle (*Mlc2*^139^ and *Act57B*^140^). For testis cell type annotation, testis bins were extracted and integrated with our adult *Dpse* testis single-cell data following Seurat label transfer workflow. Each bin was assigned to the cell type with the highest prediction score.

Bulk RNA-seq data of head, 3rd larvae, stage 9 embryos, carcasses, ovaries and testes of both *Dmel* and *Dpse*, as well as testis RNA-seq from *Dmel achi*, *mip40* and *kmg* loss-of-function (LOF) experiments were downloaded from NCBI. We used Hisat2 (version 2.0.4) ^141^ to align total RNA reads to their respective reference genomes. Gene-level read counts were obtained with FeatureCounts (version 1.6.2) ^142^. FPKM (Fragments Per Kilobase of transcript per Million mapped reads) values were calculated and used to calculate testis specificity scores as the ratio of testis FPKM to male carcass FPKM (T/C). For LOF transcriptome analysis, genes with FPKM < 1 in both LOF and wild-type (WT) samples were excluded. For the remaining genes, log2 fold change (log2FC) was calculated and genes with log2FC < –2 or log2FC > 2 were defined as *achi/mip40/kmg* regulated genes. To calculate TAU-score, customized Rcode from “severinEvo” (https://github.com/severinEvo/gene_expression) was used across RNA-seq data from stage 9 embryos, 3rd instar larvae, head, testis and ovary.

### Cell type annotation and velocity analyses

After integration, FindNeighbors and FindClusters (Seurat) were performed using 30 PCs with a resolution of 2. We annotated a total 19 different cell types, including 15 germ cell types and 4 somatic cell types. Cell identities were inferred based on established marker genes.For the germ cell lineage, cell clusters enriched for expressing *AGO3*^143^, *aub*^144^, *bam*^145^ and *vasa*^146^ were annotated as germline stem cells and early spermatogonia (G1). Cell clusters biasedly enriched for *vasa* but lacking *bam* were classified as late spermatogonia (G2). Spermatocyte populations were divided into three stages: clusters biasedly expressing *CycB*^147^ and *can, aly* and *sa*^42,43^ were annotated as early spermatocytes (C1), clusters expressing *CycB* but decreased *aly*, *can* and *sa* were annotated as middle-stage spermatocytes (C2), and those enriched for *twe*^148^, but with absent expression of *can* and *sa* were annotated as late spermatocytes (C3). Early spermatids (S1) were identified by *soti*^117^ and *bol*^149^ biased expression, while late spermatids (S2) were characterized by enrichment of *f-cup* and *wa-cup* expression^117^. For the somatic lineage, early cyst cells (Cys1) were labeled by erichment of *tj*^150^ and but not *eya*^151^ expression, whereas late cyst cells (Cys2) were labeled by enrichment of eya but not tj. Hub cells were marked by *Fas3*^152^. The cell clusters expressing *MtnA*^153^ but not *Fas3* were terminal epithelial cells (Epi).

Loom files for each sample were produced with run10X from velocyto.py (version 0.17) following the standard workflow^154^. RNA velocity analysis was then performed with scVelo (version 0.2.3) ^155^. Genes with fewer than 30 total counts were filtered out, and the top 2,000 most variable genes were kept for normalization and log-transformation. We selected the top 30 principal components and used 30 neighbors to estimate RNA velocities under the stochastic model. Velocities were projected onto the UMAP embedding generated from Seurat.

### Testis immunostaining and ChIP-seq experiments

Immunostaining was modified from ^156^. Dissected testes were torn open on a coverslip in 2-3 drops of PBS using forceps. A glass microscope slide was placed over the coverslip to squash the testes followed by snap-freezing in liquid nitrogen. The coverslip was removed and the slides were immersed into 95% ice-cold ethanol at -20 L for 10 min. Slides were then fixed with 4% formaldehyde for 7 min, washed 3 times with 0.1% PBST(PBS+0.1% Triton X-100) for 15 min each and blocked with 10% Normal Goat Serum (NGS) + PBS for 45 min. Samples were incubated 1h in room temperature with primary antibodies (diluted in 10% NGS + PBS), including 1: 500 rabbit anti-sowi (generated from Huabio), 1:1000 mouse anti-alpha Tubulin (Sigma MABS276), 1:50 mouse anti-tj (NIG-FLY 10009Ab-1), 1:1000 rabbit anti-H4K16ac (abcam ab109463), 1:200 rabbit anti-H4K20me3 (abcam ab9053), 1:1000 rabbit anti-H3K9me3 (abcam ab8898), 1:100 rabbit anti-H3K9me2 (Proteintech 39754) and 1:1000 guinea pig anti-MSL2 (shared by Regnard lab/Becker lab). Slides were then washed 3 times with 0.1% PBST for 15 min each, followed by incubation for 1h in room temperature with secondary antibodies (diluted in 10% NGS + PBS), including 1: 200 Alexa Fluor 488 conjugated anti-mouse (abcam ab150113) and 1:200 Alexa Fluor 647 conjugated anti-rabbit (abcam ab150079). After washing three times in 0.1% PBST for 15 min each, samples were mounted in a mounting medium containing DAPI (Abcam ab104139). Images were captured using a Zeiss LSM 710 laser scanning confocal microscope and processed with ZEISS ZEN software. Images were adjusted for brightness and contrast using linear or gamma correction applied to the entire image. No selective enhancement of image regions was performed.

ChIP-seq reads of all *Dpse* histone modification marks (IP) and input controls (IN) in adult male head and adult male testis were downloaded from NCBI PRJNA946626. Reads were mapped to the *Dpse* genome by bowtie2 (version 2.2.9) ^157^ with default parameters and duplicate alignments were marked and removed with samtools markdup (version 1.19.2) ^158^. Normalized IP/IN ratio of each bin was calculated with deepTools v3.5.1^159^ bamCompare function, with binsize of 1(--scaleFactorsMethod SES). We calculated the average IP/IN ratio for each gene by bedtools map.

### Gene traffic and meiotic X inactivation analyses

We combined both sequence homology and synteny information to identify the *Dpse* specific gene duplications. In brief, we focused on the orthogroups generated by Orthofiner^122^ where at all the outgroup species (*Dmel*, *Dvir* and *Dwil*) have a single gene copy located on the same homologous Muller element, or 2 species have a single copy located on the same Muller, and the remaining species lacks the ortholog, while *Dpse* has multiple homologous copies. Additional *Dpse* gene copies located on a different Muller element or lacking synteny with other species were defined as new duplicated genes. We excluded the orthogroups in which parental copies or new duplicated copies cannot be unambiguously assigned (e.g., tandem duplications on the same chromosomes). Similarly, in the orthogroups where *Dmel*, *Dvir* and *Dwil* have a single gene copy located on the same Muller element, but *Dpse* has a single gene copy located on a different Muller element, such *Dpse* genes were defined as translocated genes. For the gene duplications and translocations shared by *D. subobscura* were identified using the same criteria. Interchromosomal gene traffic (also referred to as relocations in the ms) include all gene translocations and gene duplications that generated only one new duplicated gene located on a different chromosome from its parental gene. Because the AX and DX arms of the *Dpse* X chromosome originated at different evolutionary time points, we applied different criteria to infer gene movement. For gene traffic involving AX (AX-to-autosome or autosome-to-AX), we used *Dmel* as an outgroup. For gene traffic involving the DX (DX-to-autosome or autosome-to-DX), we used *Dsub* and *D. bifasciata* as outgroups, ensuring that the identified traffic genes originated after the formation of the DX arm.

We inferred the traffic genes that are meiotic silenced with the following criteria: for translocations, both translocated gene and parental gene should have CPM value >0.5 at early spermatocytes (early meiotic stages) in testis scRNA-seq; for cases of duplications, new duplicated genes are expected to have higher expression (*P* <0.05, one-sided Wilcoxon tests) in early spermatocytes than that of parental genes. For demasculinization gene movements, parental genes are expected to have FPKM value > 1 in testis bulk RNA-seq, and CPM> 0.5 in testis scRNA-seq, and also meet any one of the 3 items: 1) the translocated or duplicated copy has a testis specificity (T/C) lower than than half that of parental gene; 2) the testis FPKM value of translocated or duplicated copy is lower than half the value of parental gene; 3) the average scRNA-seq CPM value of the translocated or duplicated copy is lower than half the value of parental average CPM value.

We followed the procedure described in ^53^ to analyze the interchromosomal gene movements. First, for each type of gene movement we calculated the expected ratio of movement, which was defined as the ratio between the number of genes on the source chromosome and the euchromatic genome size of the target chromosome. Relocated genes that contain only one exon whereas their parental genes containing more than one exon were defined as RNA-based gene movements (retrotransposition) and relocated genes with more than one exon were defined as DNA-based movements. The frequency of retrotransposition depends on the relative population sizes of the chromosomes, that is 0.75 when the X chromosome is the target chromosome, and 1 when the X chromosome is the source chromosome. The frequencies of DNA-based movements for the X chromosomes and autosomes are equal to 1, because males do not have crossing-over^55^. Then the expected number of gene movements were calculated based on the expected frequencies and total number of gene relocation events. The observed number of gene movements corresponded to relocated genes identified through duplication or translocation events. Statistical significance between expected and observed gene transport numbers was assessed using a chi-square test.

### Dynamic Network Biomarkers (DNB) analyses

We modified the landscape dynamic network biomarker (l-DNB) method from ^79^. We assumed each cell type contains *n* cells and *m* genes, its DNB score *I*_DNB_, is defined as the average of the top *k* gene local DNB score:

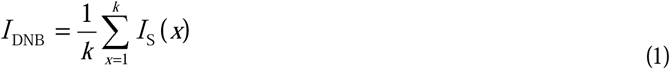

where *I*_s_(*x*) is the local DNB score of gene *x*. And the calculation of *I*_s_(*x*) includes the following five steps:

1. construct a gene correlation network. Calculate the Pearson correlation coefficient *PCC* (*x*, *y*) and its significance *P-value* between any two genes, say gene *x* and gene *y*. If and only if *P-value* < 0.01, the correlation is considered meaningful and exists, and 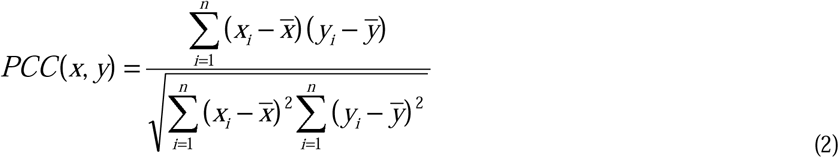
2. calculate the average standard deviation of the module where gene *x* is located. The module where gene *x* is located refers to the gene set composed of gene *x* and its first-order neighbors, and its average standard deviation is 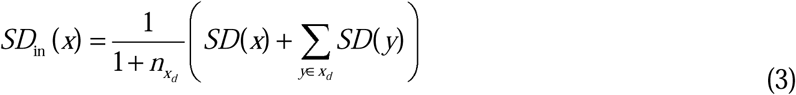 Where, *SD* (*x*) represents the standard deviation of expression of gene *x*, *x_d_* represents the first-order neighbor set of gene *x*, and 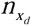 represents the number of elements in set *x_d_*.
3. calculate the average Pearson correlation coefficient within the module where gene *x* is located. Here we calculated the average absolute value of the Pearson correlation coefficient between gene *x* and its first-order neighbors, 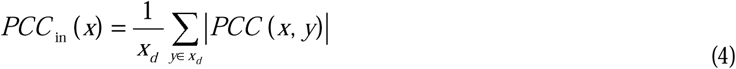
4. calculate the average Pearson correlation coefficient between the internal and external of the module where gene *x* is located. What we are calculating here is the average absolute value of the Pearson correlation coefficient between the first-order neighbors and the second-order neighbors of *x*, 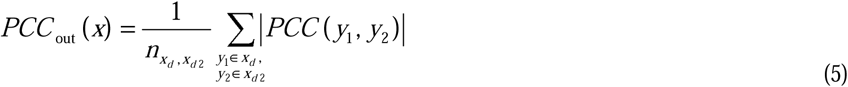
5. calculate the local DNB score of gene *x*. The local DNB score of gene *x* is defined as

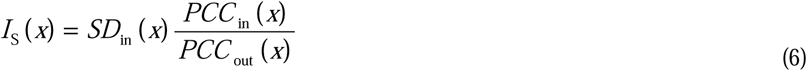

For both larva and adult, the 500 genes with the highest DNB index were defined as DNB genes (DNBGs).

### Differentially Expressed Genes (DEGs) between sperm types

Differentially expressed genes (DEGs) between eusperm and parasperm were identified using the FindMarkers function in Seurat, based on single-cell transcriptomes from both adult and larval testes. Genes were considered significantly differentially expressed if they satisfied the thresholds of log2 fold change (log2FC) > 0.5 and *q-*value < 0.05. Any gene showing differential expression between eusperm and parasperm in at least one cell type was classified as a differentially expressed gene between eusperm and parasperm (EP-DEG). Using the same criteria, we also identified DEGs between the two parasperm subtypes (PP-DEG). Gradient genes were defined as those exhibiting gradient expression differences among the three sperm types (either eusperm > parasperm2 > parasperm1 or eusperm < parasperm2 < parasperm1). To be classified as a gradient gene, a gene was required to have a log2 fold change (log2FC) > 0.25 between each pair of adjacent sperm types, with at least one comparison showing log2FC > 0.5. The same selection criteria were applied to identify gradient genes for all possible permutations of the three sperm types. These two ordered gradient patterns (eusperm > parasperm2 > parasperm1 and eusperm < parasperm2 < parasperm1) were significantly overrepresented compared with all possible permutations (Chi-square test, *P* < 0.001).

Gene ontology enrichment for DNBGs, DEGs and gradient genes were performed using Metascape^160^.

### Predominantly transcribed testis cell types for each gene

We calculated the average expression of each gene across germ cell stages, including germline stem cell and early spermatogonia (G1), late spermatogonia (G2), early-to-middle spermatocytes (C1-C2), late spermatocytes (C3), early spermatids (S1) and late spermatids (S2) using the AverageExpression function (Seurat v4). Genes with a maximum average expression less than 0.2 across cell types were excluded. Expression values were subsequently normalized by z-score transformation. Gene expression heatmaps were generated using the ComplexHeatmap R package^161^. Genes were grouped into 10 clusters using the row_km option in Heatmap() and cluster membership was extracted for downstream analysis. Based on the highest expression stage, genes were assigned to 5 stages: spermatogonia (G1-G2), late spermatogonia and early-to-middle spermatocytes (G2-C2), spermatocytes (C1-C3), late spermatocytes and early spermatids (C3-S1), late spermatids(S2) (**Supplementary Fig. 24**).

We applied gene expression stages to investigate functional divergence among new duplicated genes in *Dpse*. The *Dmel* ortholog was treated as representing the ancestral expression pattern. For each gene duplication pair, we compared the expression stage of both new duplicated copy and parental copy to that of the *Dmel* ortholog. Gene duplication pairs were classified into three functional categories: 1) silenced-in-testis, where either of new copy or the parental copy showed no detectable expression while the other retained the same expression as *Dmel* gene; 2) conserved-cell-types, where both new and parental copies exhibited expression consistent with the *Dmel* gene; 3) neo-functionalization, where at least one copy exhibited a novel expression stage distinct from the *Dmel* gene. Gene pairs were also defined as neo-functionalization if either the new copy or parental copy gained their testis-specific expression pattern. In detail, one of the *Dpse* copies should fulfill all of the following criteria: 1) high testis expression (FPKM >1); 2) strong testis specificity (T/C >2); 3) lack of strong testis specificity in the *Dmel* ortholog (T/C<2), and (iv) a ≥4-fold increase in testis specificity relative to *Dmel* ortholog. Duplications in which neither the new copy nor the parental copy nor the *Dmel* ortholog were expressed were excluded. For single-copy genes, functional categories were defined analogously by comparing the expression stages of orthologous genes between *Dpse* and *Dmel*. Three functional classes were assigned based on whether the expression stage of the *Dpse* gene matched or diverged from that of its *Dmel* ortholog, or whether the *Dpse* gene gained testis specificity.

To calculate dN/dS value protein sequences of single copy orthologous genes from 4 *Drosophila* species (*D. melanogaster*, *D. pseudoobscura*, *D. virilis*, and *D. willistoni*) were aligned using PRANK v150803^162^. Conserved alignment blocks were extracted with Gblocks v9.1b^163^, and the resulting alignments were converted into PHYLIP format. To estimate evolutionary rates, we ran codeml in the free-ratio model implemented in PAML v4.8^164^, which provides branch-specific estimates of the ratio of nonsynonymous to synonymous substitution rates (dN/dS). For downstream analyses, genes with extreme or unreliable values (dS < 0.01 or dS > 1) were excluded.

## Data availability

Raw reads of scRNA-seq and stereo-seq are available on NCBI Sequence Read Archive (SRA) under accession number PRJNA1271262. The *D.pse* ChIP-seq datasets used in this study are available from the NCBI under BioProject accession PRJNA946626. The *D.mel achi/vis* and *kmg* loss of function data are available from the NCBI under BioProject accession PRJNA450930 and PRJNA352407.

## Code availability

All codes developed in this work is available at https://github.com/hehuangyi/Fly-single-cell.

## Acknowledgments

We thank Rhonda Snook and Helen White-Cooper for their very helpful comments throughout the manuscript. This work was funded by the National Key Research and Development Program of China (2024YFA1802500 and 2023YFA1800500), and National Natural Science Foundation of China (73432170415) to Q.Z.

## Author contributions

Q.Z. conceived the project; H.H., L.C., M.A., L.Y. collected the data and performed the experiments; H.H., Q.Z., Q.S., J.J., M.W., R.X., X.X., L.C. analysed the data; H.H. and Q.Z. wrote the manuscript.

