## Supplementary Figures for "New genes play a prominent role in evolution of new sperm classes of *Drosophila*"

### Fig. S1

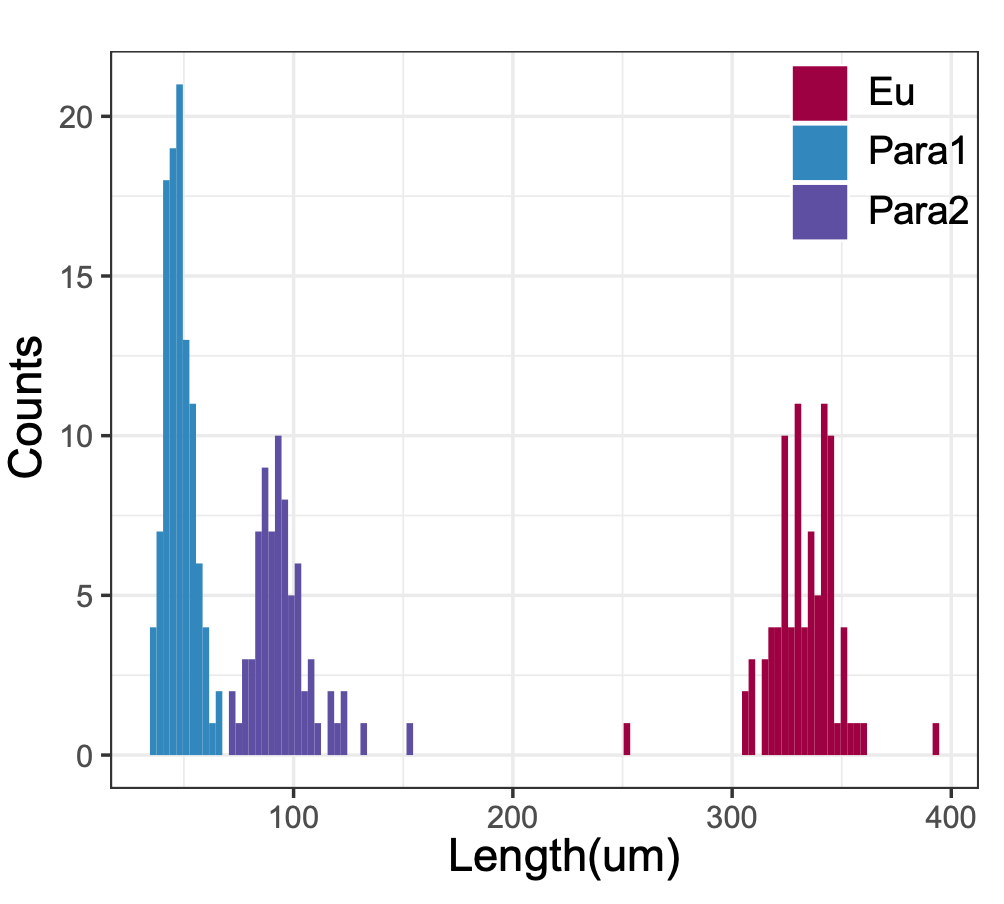

**Fig. S1 Sperm length distribution in *Dpse*.** Sperms collected from seminal vesicles were distributed into three non-overlapping categories by their lengths: longest eusperm (330.17 μm ± 28.86 SD), shortest parasperm1 (48 μm ± 7.49 SD) and mid-short parasperm2 (95.96 μm ± 13.27 SD).

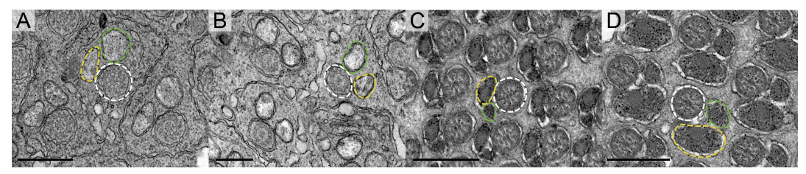

**Fig. S2 Transmission Electron Microscopy (TEM) images of *Dpse* sperm tails.** Figure shows the ultrastructure of spermatids at the mid-elongation (A, B) and late-elongation (C, D) stages of *Dpse*. Each spermatid contains one axoneme (white dashed line), one major mitochondrial derivative (yellow dashed line) and one major mitochondria derivative (greed dashed line). At the same developmental stage, spermatids showed no detectable ultrastructural differences. This is consistent with what previously found in *Drosophila subobscura*, that the eusperm and parasperm have similar architecture *(127)*. Scale bar: 0.5 μm.

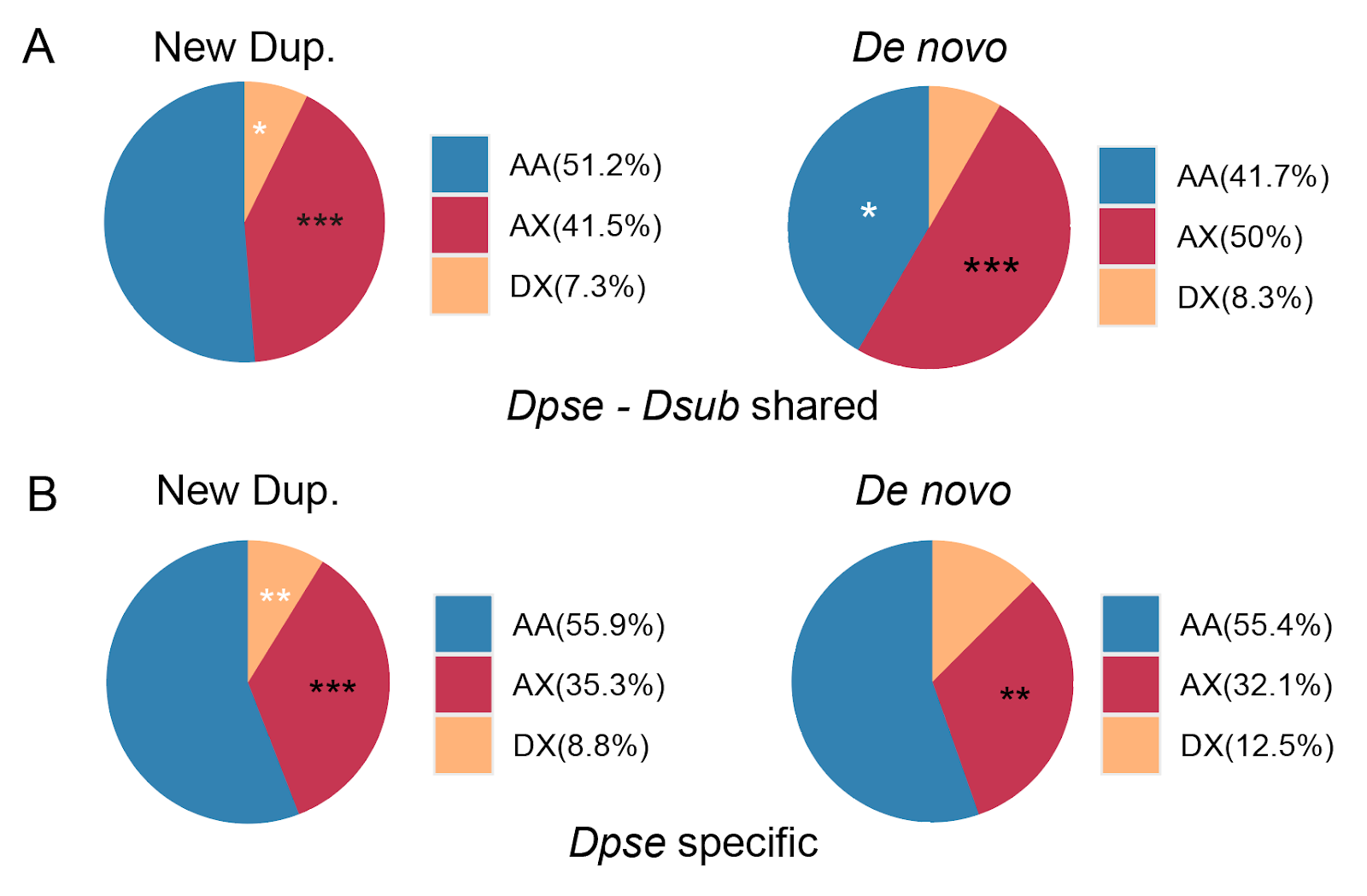

**Fig. S3 Chromosome distribution of new duplicated genes and *de novo* genes in *Dpse.*** Compared with single copy genes shared in *Dmel*, *Dpse,* and *Dsub*, both duplicated genes and *de novo* genes in *Dpse* are significantly enriched on the AX chromosome. This enrichment is observed for new genes that originated from the ancestor of the obscura group (A; shared by *Dpse* and *Dsub*) as well as for genes that arose after species divergence and sex chromosome turnovers in *Dpse* (B; *Dpse*-specific). Autosomes (AA), ancestral X (AX), and new-X (DX) are indicated. Black asterisks denote significant enrichment (*P* < 0.05, **P* < 0.01, ***P* < 0.001), whereas white asterisks denote significant depletion.

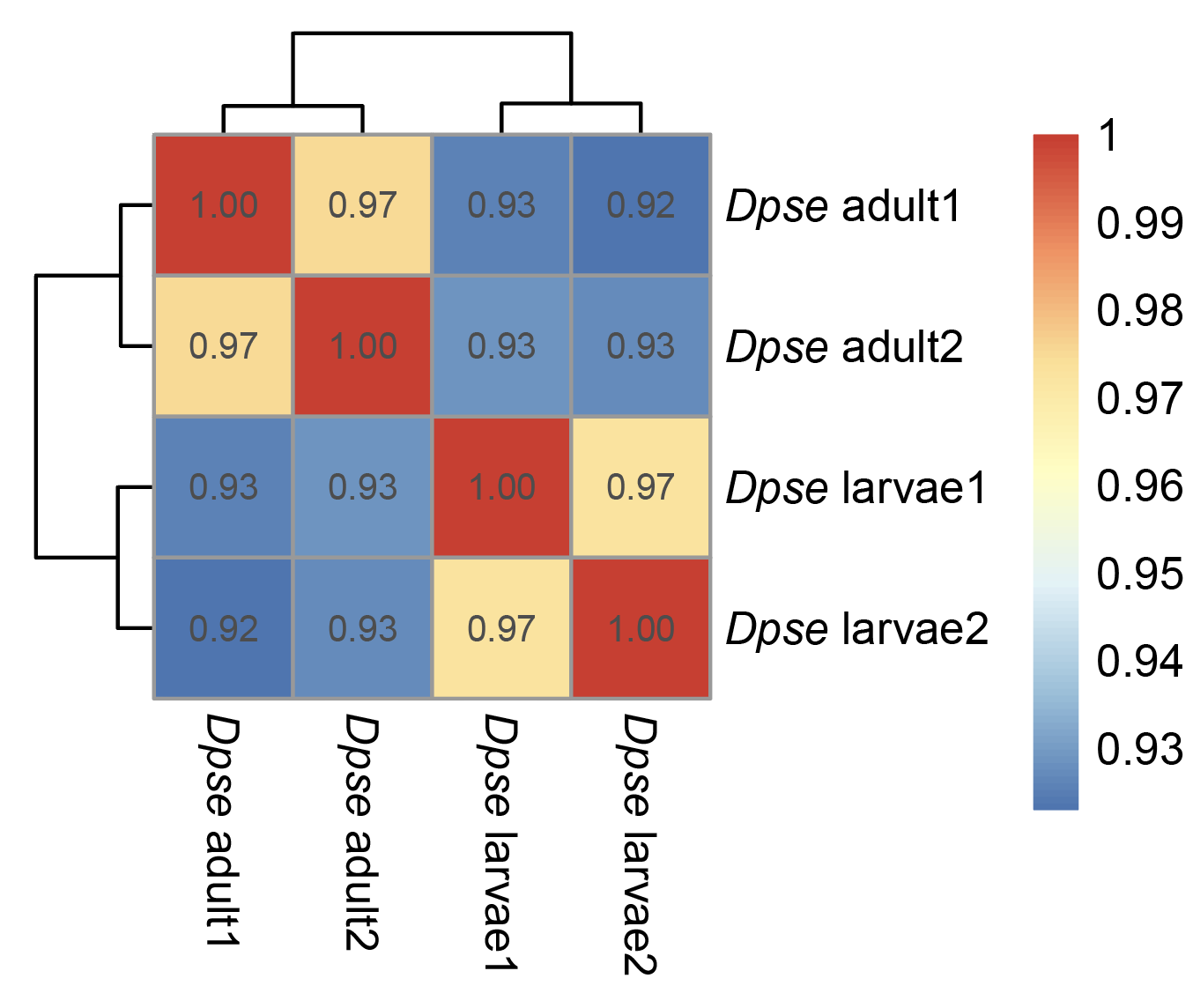

**Fig. S4 Heatmap of Pearson correlation coefficients between Dpse testes scRNA-seq samples.**The heatmap shows pairwise Pearson correlation values among four *Dpse* testes scRNA-seq samples using total UMI for each gene. Correlation coefficients range from 0.93 to 0.97, indicating high similarity in gene expression profiles across samples.

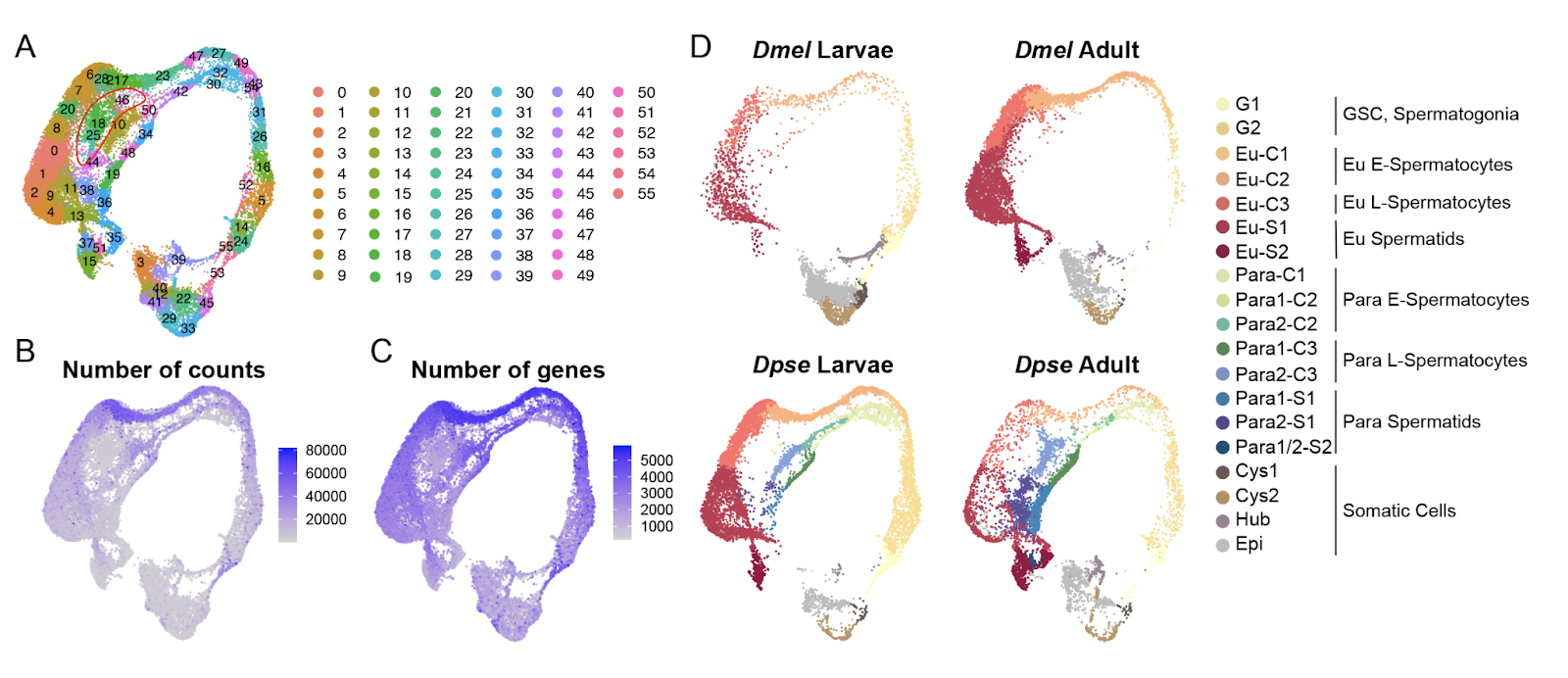

**Fig. S5 Uniform Manifold Approximation and Projection (UMAP) of single-cell transcriptomes from *Drosophila* testes across species and developmental stages.** (A) UMAP embeddings of integrated *Dpse* and *Dmel* testes cells, clustered using the FindClusters function. Clusters outlined in red show low numbers of transcripts. (B) and low numbers of detected genes (C) and were therefore excluded from downstream analyses. (D) UMAP embeddings of testes cells grouped by species (*Dmel* and *Dpse*) and developmental stage (larva and adult). Cell type annotations are indicated on the right.

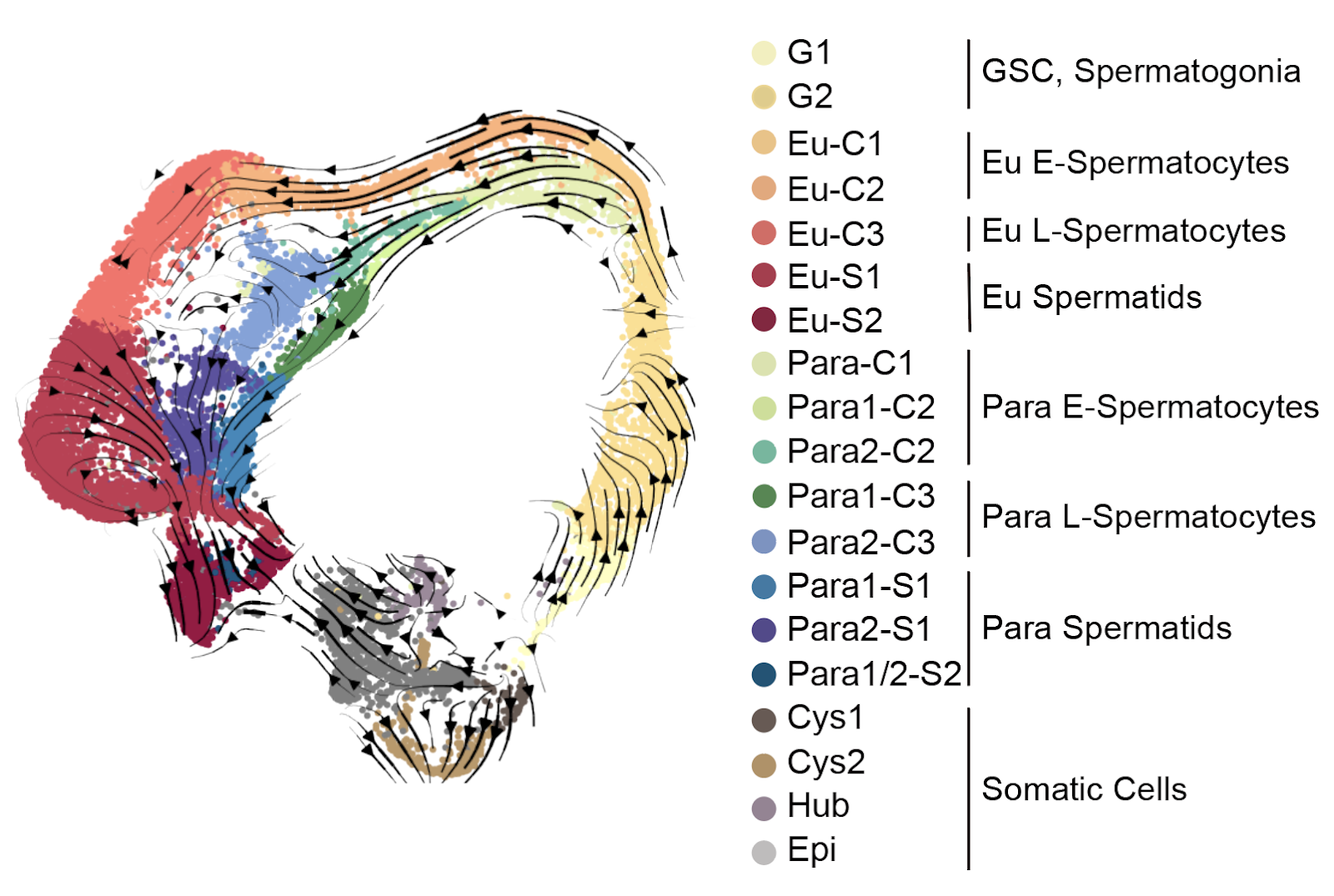

**Fig. S6 RNA velocity analysis of *Dpse* testes single-cell transcriptomes.** The flow lines indicate the directionality of cell state transitions inferred by RNA velocity.

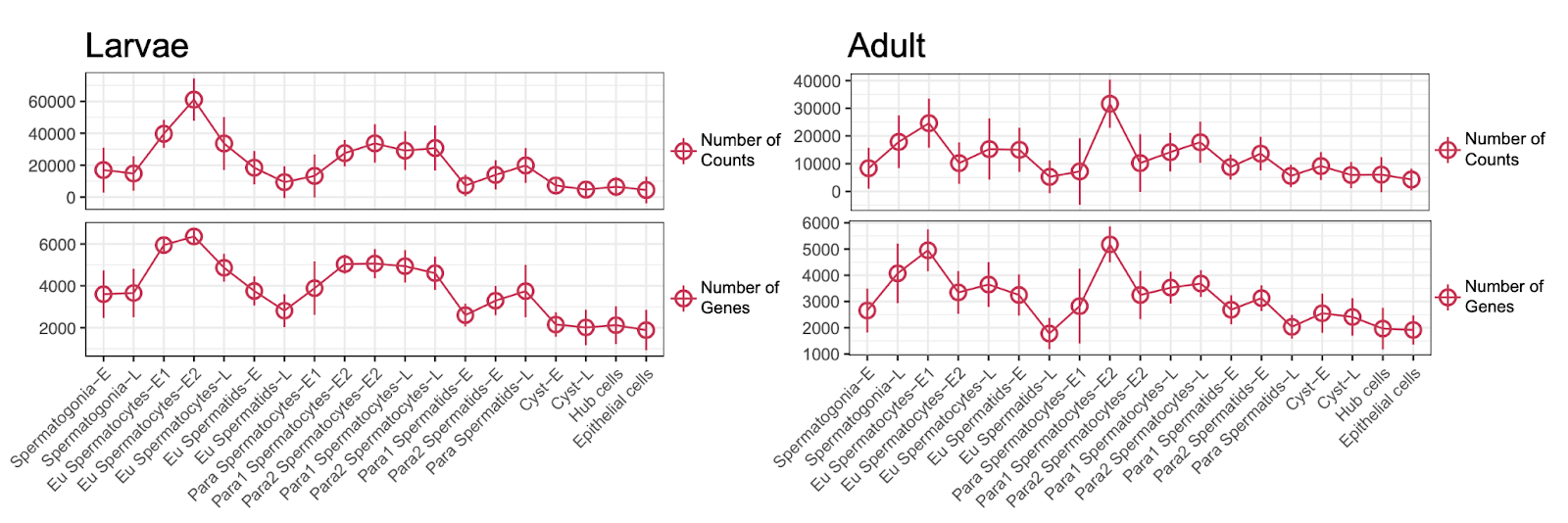

**Fig. S7 number of UMI counts and number of expressed genes for each cell type in adult and larvae.**

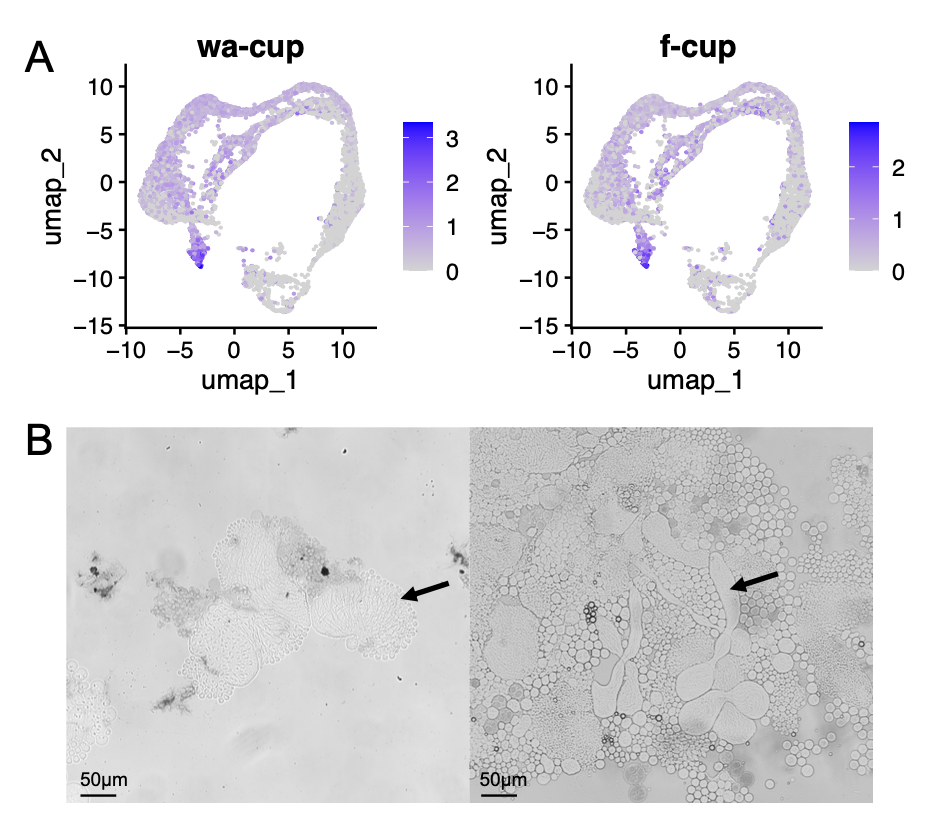

**Fig. S8 Identification of late spermatids in *Dpse* larval testes.** (A) Expression of late spermatids markers *wa-cup* and *f-cup* in *Dpse* larval testes scRNA-seq data. (B) Elongating spermatids (arrow) observed in third-instar larval testes under an optical microscope.

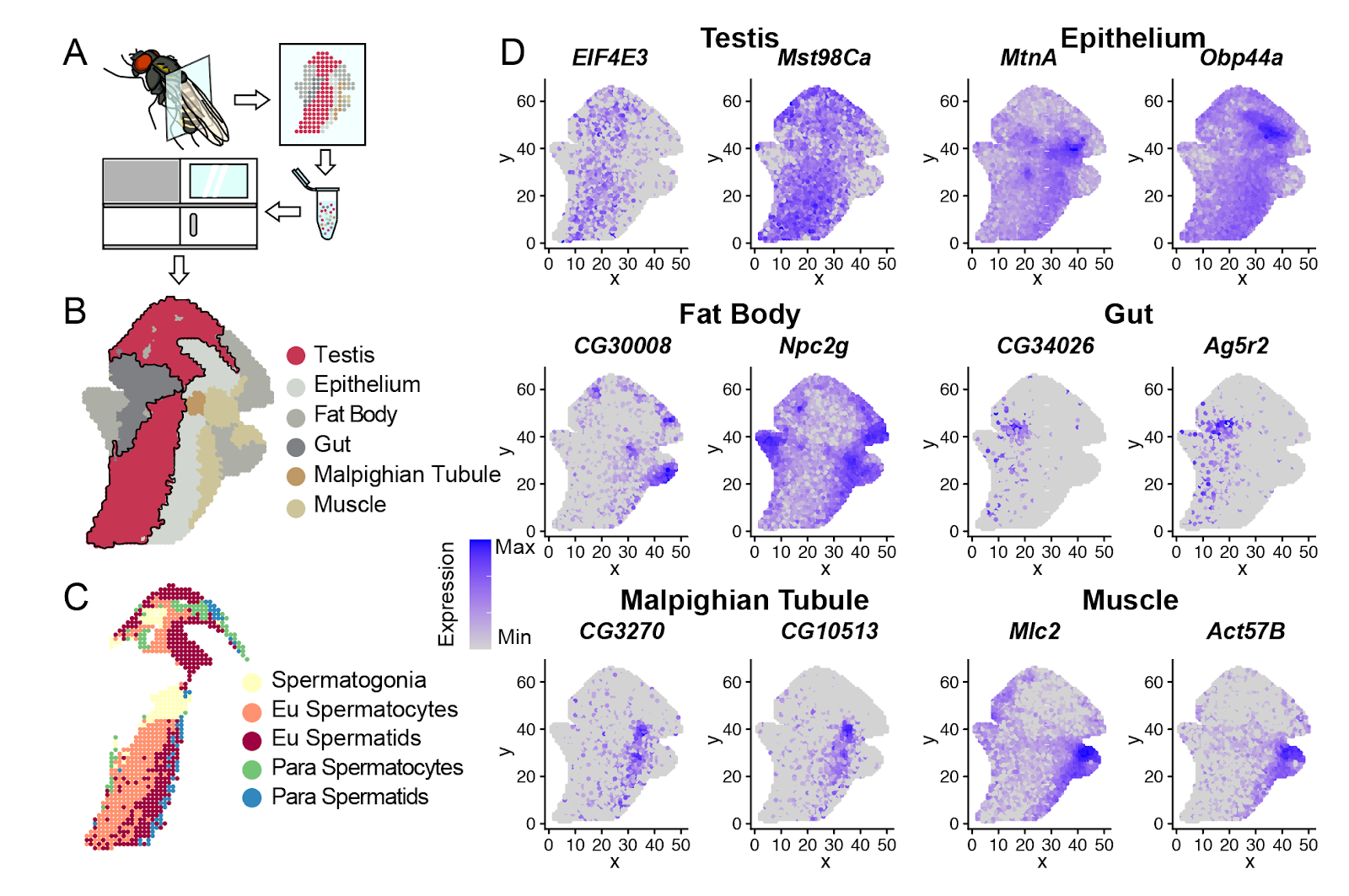

**Fig. S9 Spatial transcriptomic mapping of *Dpse* male testis and somatic tissues.** (A) Workflow of spatial transcriptomic profiling from adult male *Dpse* abdomen part. (B) Spatial annotation of major tissue types, including testis, epithelium, fat body, gut, Malpighian tubule, and muscle. (C) Fine-scale annotation of testis subregions, highlighting spermatogonia, eusperm spermatocytes, eusperm spermatids, parasperm spermatocytes, and parasperm spermatids. (D) Spatial expression patterns of representative marker genes for each tissue type. Color scale represents normalized expression levels from low (gray) to high (blue).

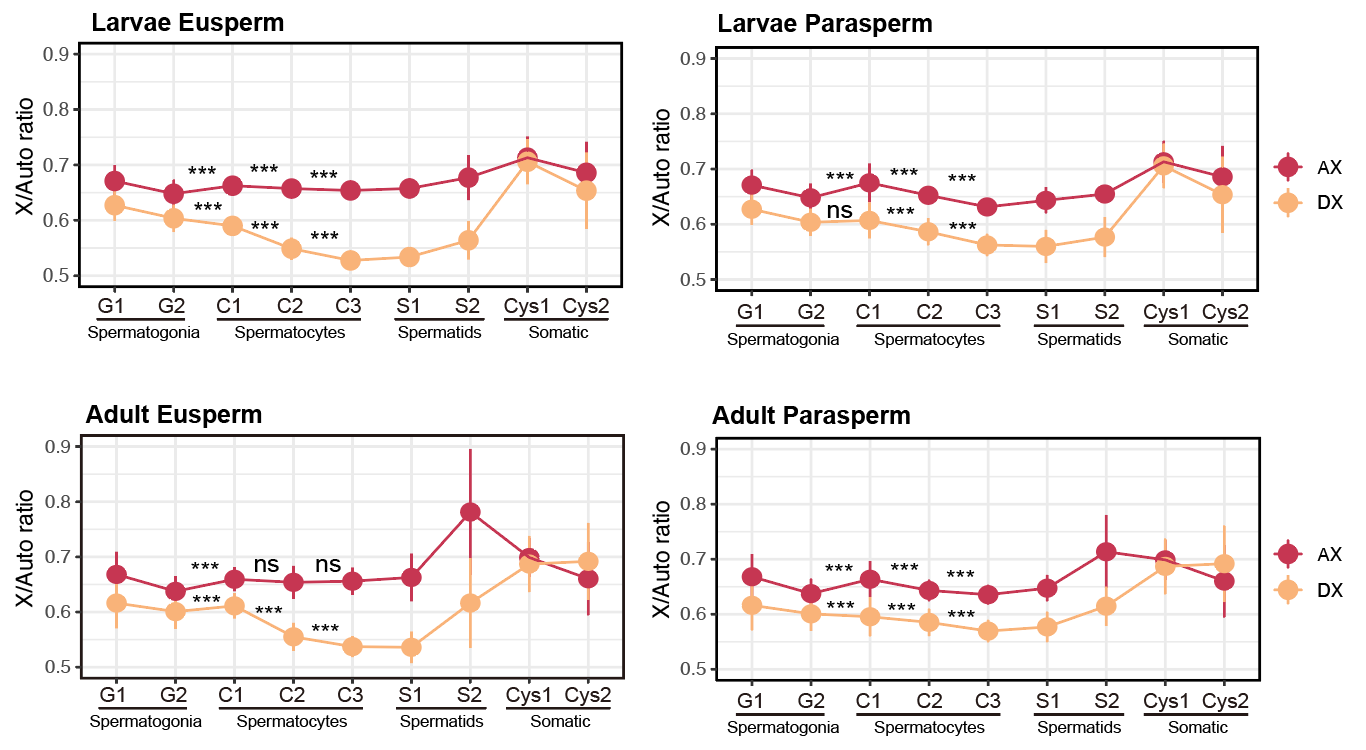

**Fig. S10 Relative expression of X to Autosomes across cell types in adult eusperm, adult parasperm, larval eusperm and larval parasperm.** Y-axis is total counts of genes from AX or DX chromosomes divided by that from autosomes (Muller B, C and E) normalized by the number of genes per chromosome. Each dot represents the median value of X to Autosome ratios of each cell type. The error bar represents the standard deviation of each cell type relative to the median value. Comparison expression ratio in adjacent cell types were tested by Wilcoxon one-side, *** *P* < 0.001.

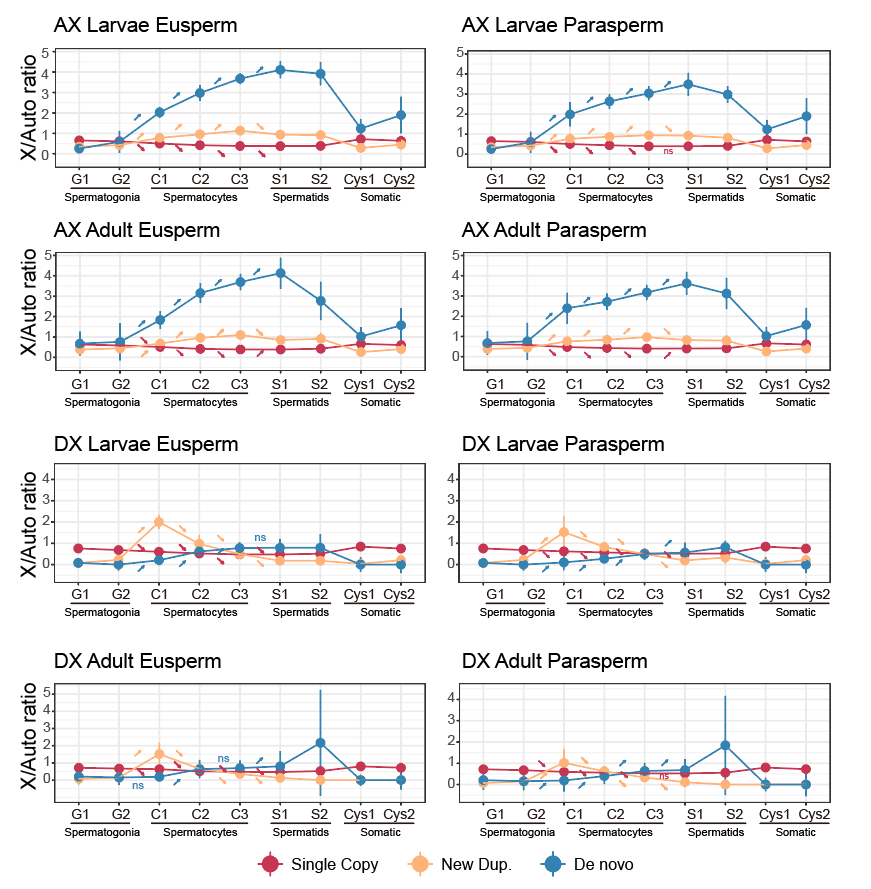

**Fig. S11 Relative expression of X-linked single copy gene, new duplicated genes and de novo genes.** Y-axis is total counts of genes from X-lined single copy genes, new duplicated genes or de novo genes divided by that from autosomes (Muller B, C and E) normalized by the number of genes. Each dot represents the median value of gene to autosome ratios of each cell type. The error bar represents the standard deviation of each cell type relative to the median value. Arrows show the ratio of the later stage is significant (p < 0.001, One-sided Wilcoxon test) higher or lower than the former stage.

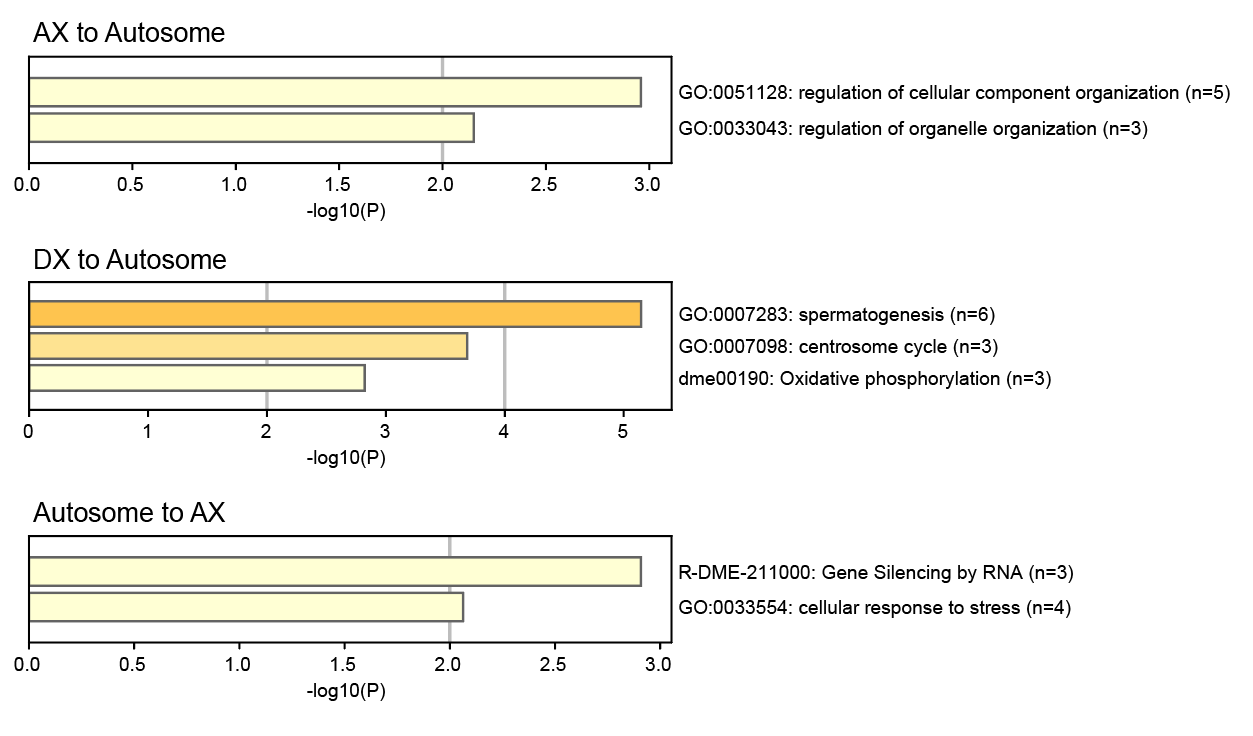

**Fig. S12 Gene Ontology (GO) enrichment of genes involved in different chromosomal relocation events.** Bar plots show significantly enriched functional categories for genes relocated from the AX to autosomes (top), from the DX to autosomes (middle), and from autosomes to the AX chromosome (bottom). Bar length represents the statistical significance (−log10 *P* value), and GO terms or pathways are listed to the right of each plot.

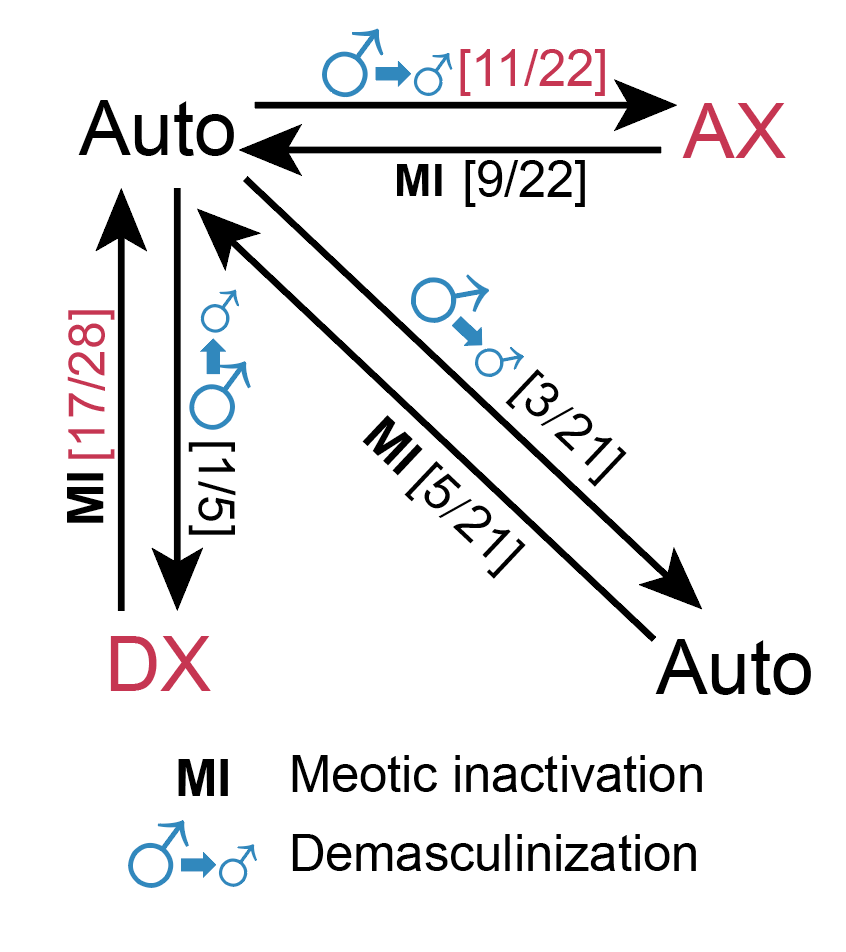

**Fig. S13 Gene relocation associated with meiotic inactivation (MI) and demasculinization between X chromosomes and autosomes.** Arrows indicate gene relocation events from the source chromosome to the target chromosome. The number of gene relocation events that meet the criteria for meiotic inactivation and demasculinization, and the total number of relocation events between the two chromosomes are shown in the square brackets. Compared to autosome-to-autosome events, statistically significant categories are highlighted in red (*P* < 0.05, one-sided Fisher’s exact test**).**

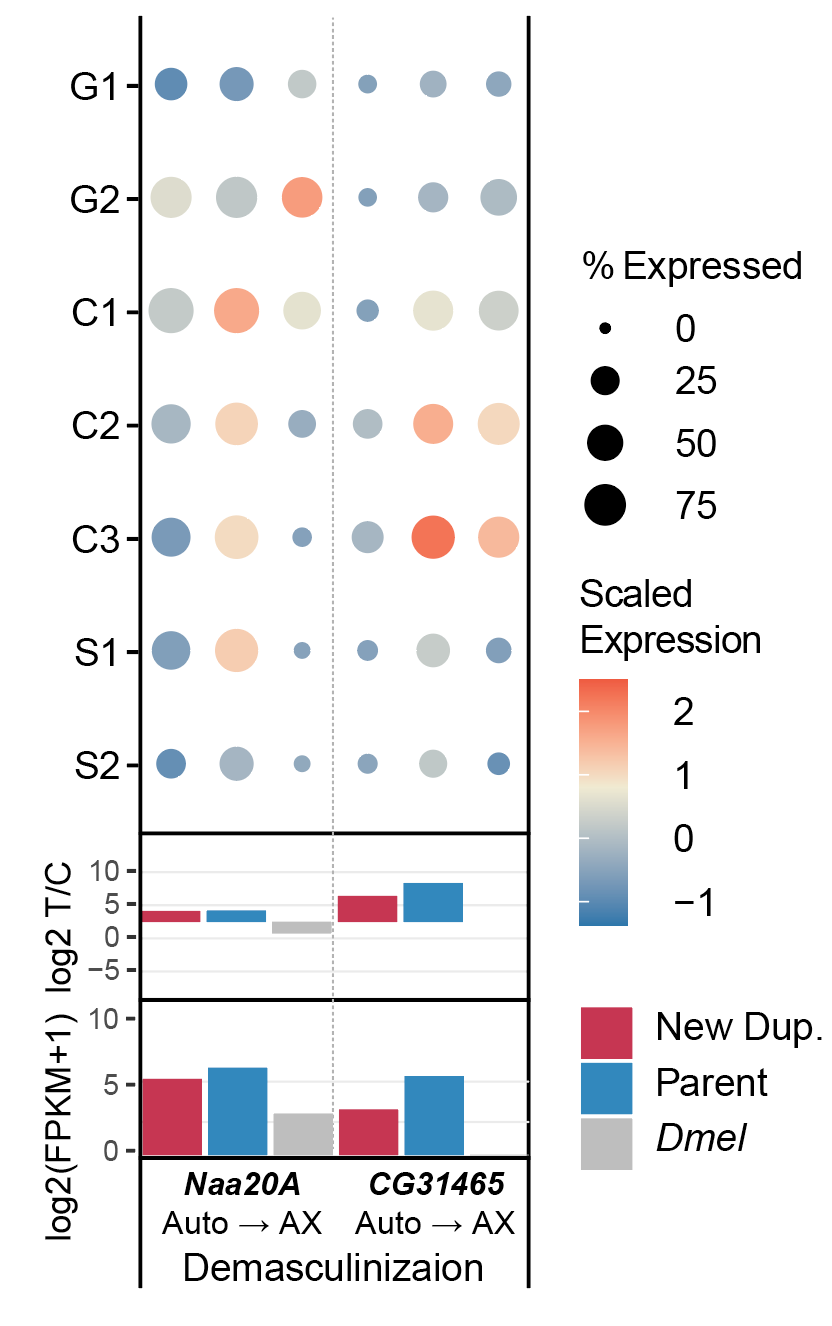

**Fig. S14 Expression patterns of relocated genes during *Drosophila* spermatogenesis.** Single cell and bulk RNA-seq expression profile of gene relocation cases supporting demasculinization. The circle size is scaled to the percentage of cells with expression of each marker gene and the color is scaled within the same species for each gene group. The spermatocytes (C1-C3) and spermatids (S1, S2) data is from eusperm. Red: new duplicated genes; blue: parental gene; grey: *Dmel* orthologous.

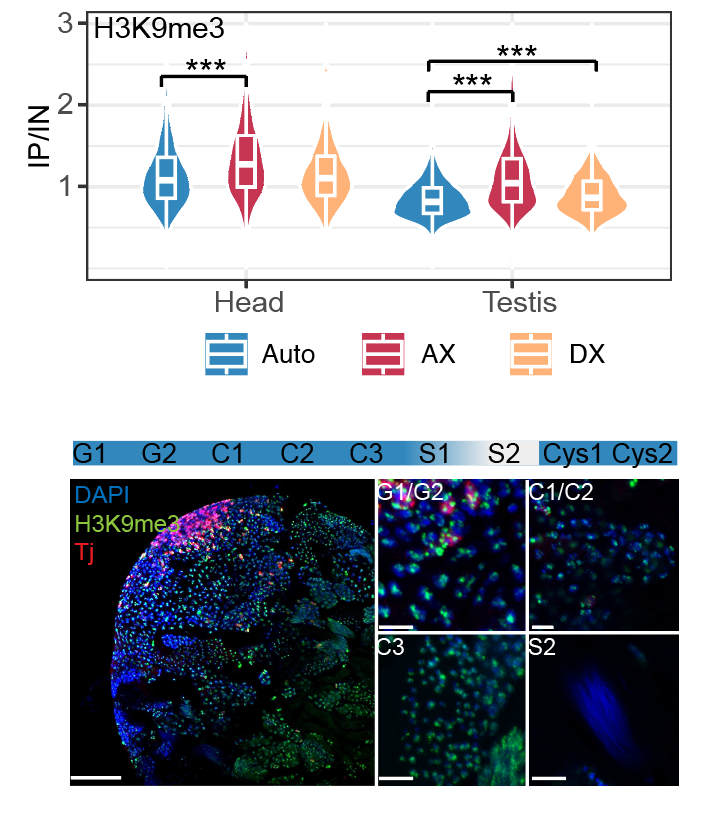

**Fig. S15 Testis enrichment and immunofluorescence of H3K9me3.** (A)H3K9me3 enrichment (IP/Input) of genes in heads and testes from autosomes (Auto), AX, and DX.Boxes indicate the median and interquartile range within violin plots; ****P* < 0.001(Wilcoxon two-sided). (B) Immunofluorescence staining in the testis. Panels show the whole-mount apical region of the testis (left), spermatogonia (G1/G2) and early spermatocytes (C1/C2), late spermatocytes (C3), and late spermatids (S2) (right). Blue, DAPI; green, anti-H3K9me3; red, anti-Traffic jam (Tj), which labels early cyst cells. Scale bars, 50 µm (left) and 10 µm (right).

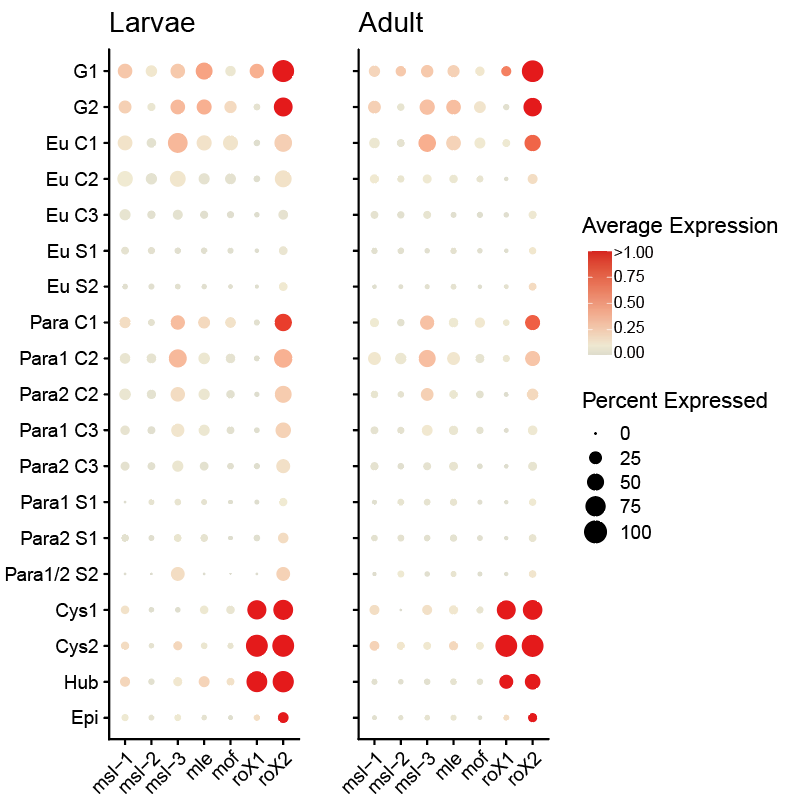

**Fig. S16 Expression patterns of male-specific lethal (MSL) complex genes in single-cell transcriptomes of larval (left) and adult (right) testes.** Dot size represents the percentage of cells expressing each MSL complex gene, and dot color indicates the average expression level within each cell type.

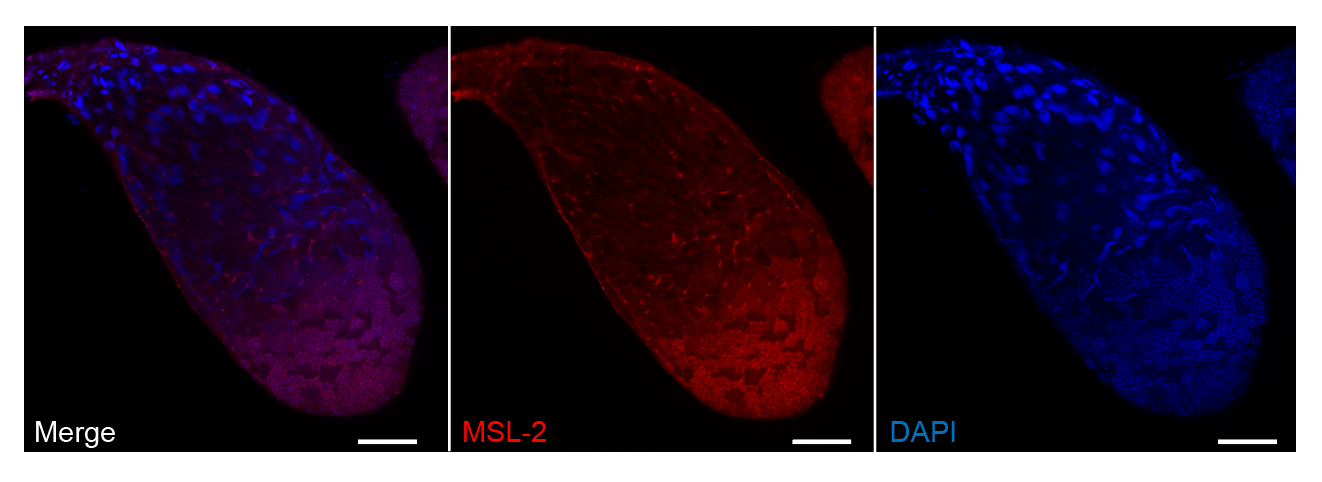

**Fig. S17 Immunofluorescence staining of Msl-2 in testis.** Blue, DAPI; Red, anti-MSL-2; Scale bars, 100 µm.

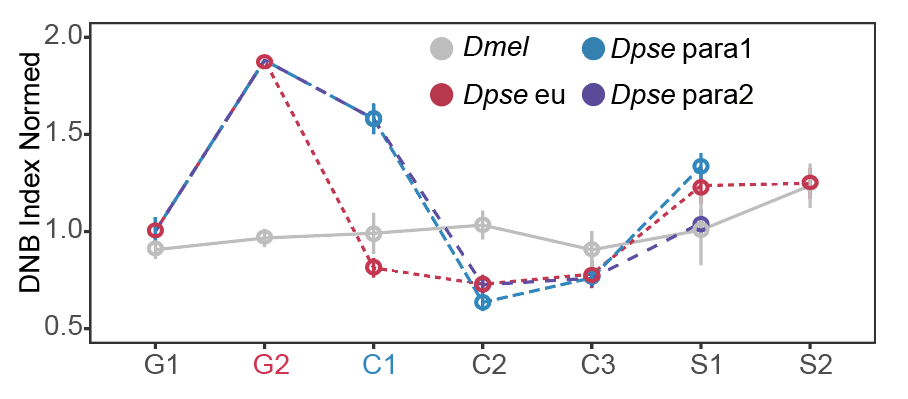

**Fig. S18 Normalized DNB index at each cell type for larva data.** Dot shows the average DNB index of the top 500 genes normalized by the average DNB index of all genes. The error bar represents the standard deviation relative to the mean value.

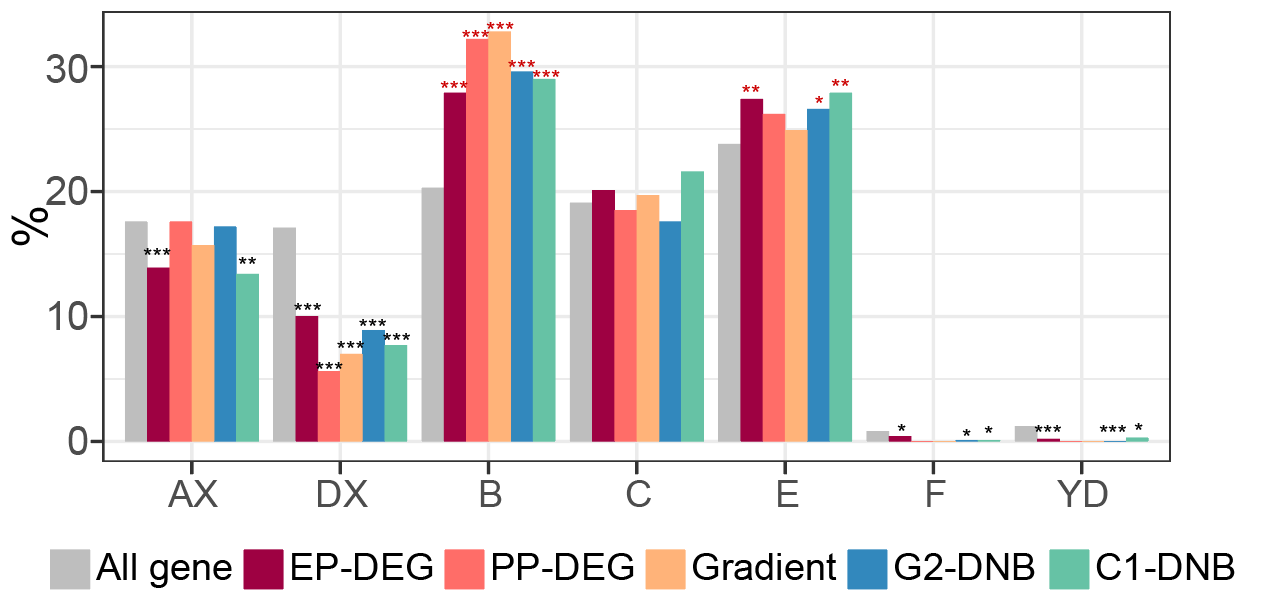

**Fig. S19 Distribution of different gene sets across Muller elements.** Bar plots show the percentage of genes located on each Muller chromosome (AX, DX, B, C, E, F, and YD) for all genes (gray) and for different gene sets, including EP-DEGs, PP-DEGs, gradient genes, G2-DNBs, and C1-DNBs. Asterisks denote significant deviations in chromosomal distribution compared with all genes. Red and black asterisks denote significant enrichment and depletion, respectively. * *P* < 0.05, ** *P* < 0.01, *** *P* < 0.001 (Fisher-exact test).

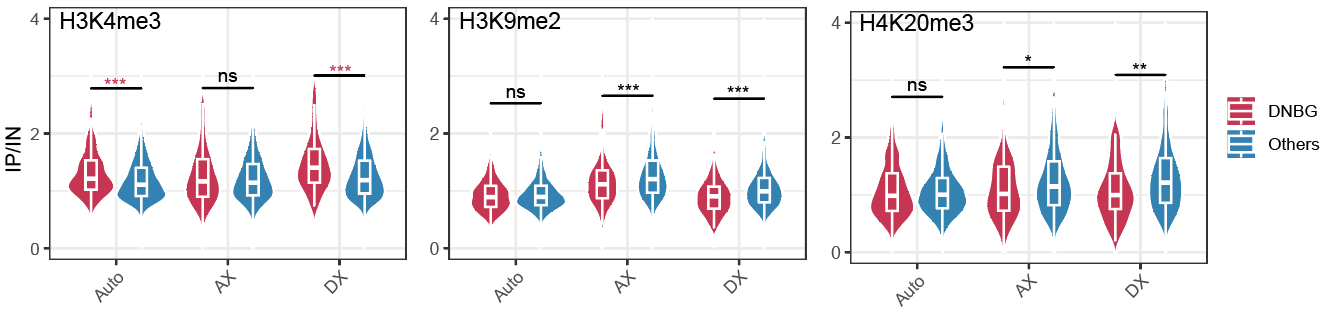

**Fig. S20 Comparisons of histone modification enrichment between C1-DNBGs and other genes.** Violin plots show the distribution of ChIP signal enrichment (IP/IN) for H3K4me3, H3K9me2, and H4K20me3 across autosomes (Auto), AX, and DX. Boxplots embedded within violins indicate the median and interquartile range. Statistical significance between DNBGs and other genes was assessed using the Wilcoxon test. * *P* < 0.05; ** *P* < 0.01; *** *P* < 0.001.

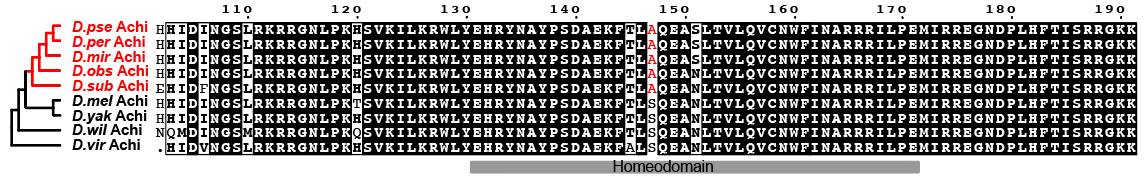

**Fig. S21 Sequence alignment of par of Achi proteins across *Drosophila* species.** The amino acid positions indicated at the top are based on the *Dpse* sequence. The conserved homeodomain is highlighted by the gray bar. The phylogenetic relationships among Achi orthologs are shown on the left. Species highlighted in red belong to the obscura group. Conserved residues are shown in black, and gaps are indicated by dashes. The red amino acid indicates *obscura*-group–specific substitution in the Achi proteins.

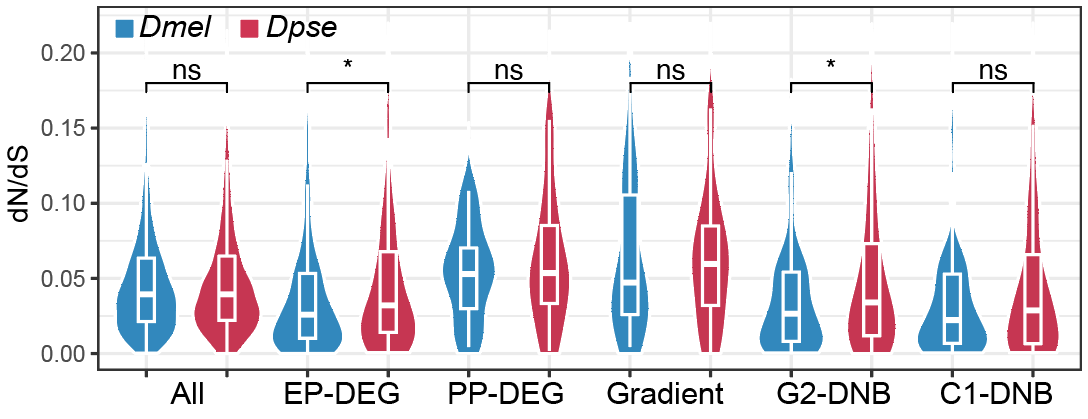

**Fig. S22 Comparison of nonsynonymous to synonymous substitution rate (dN/dS) between *Dmel* (blue) and *Dpse* (red) orthologous genes.** Violin plots show the dN/dS value for all genes, EP-DEGs, PP-DEGs, gradient genes, G2-DNBs, and C1-DNBs. Boxplots embedded within violins indicate the median and interquartile range. Statistical significance was assessed using the two-sided Wilcoxon test. * *P* < 0.05.

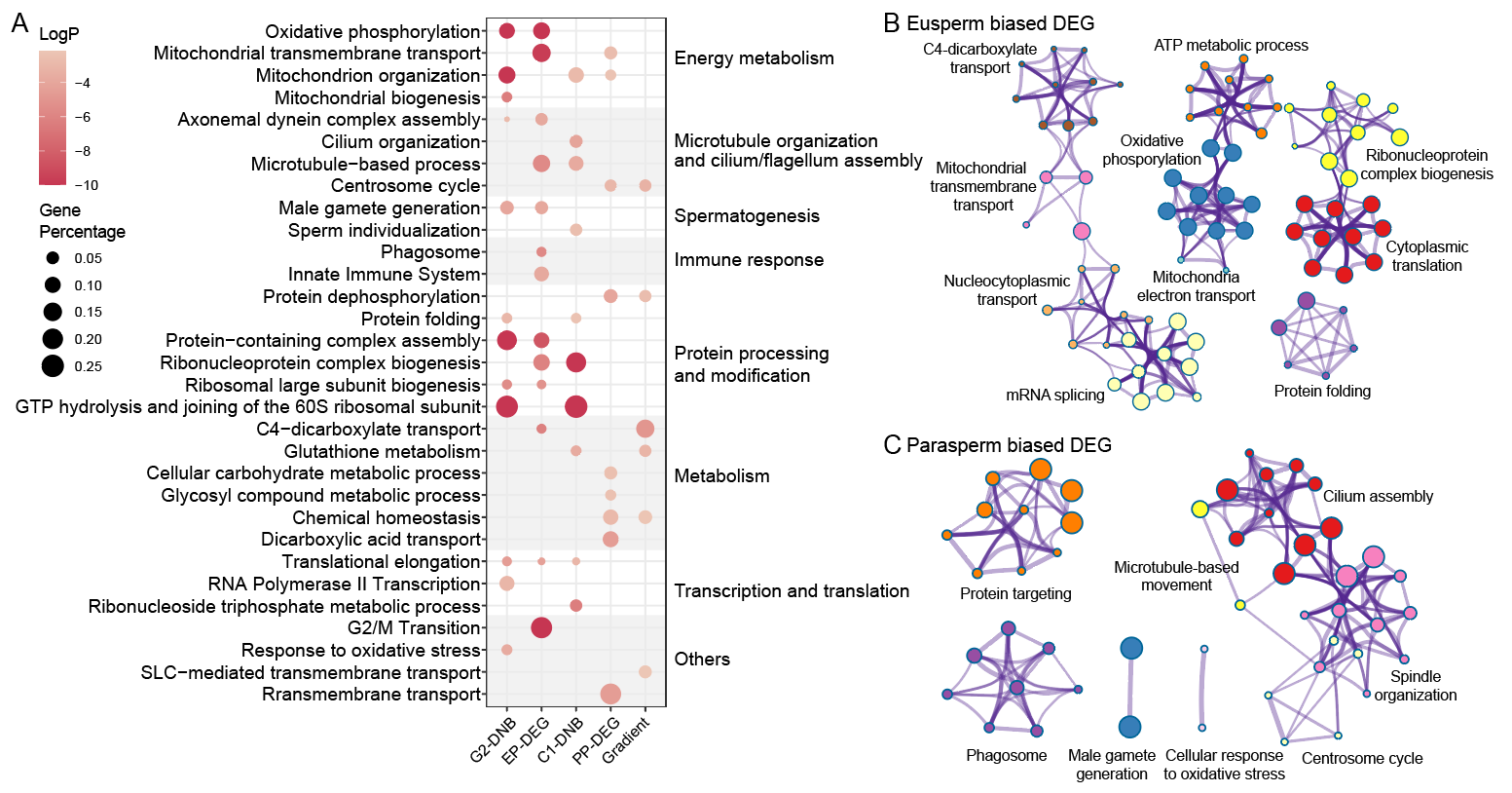
 **Fig. S23 Gene Ontology (GO) enrichment across different gene sets.** (A) Bubble plot of GO enrichment across gene sets. The x-axis represents different gene sets and the y-axis denotes GO terms. Bubble size is proportional to the number of enriched genes, and color reflects the enrichment significance. (B-C) GO enrichment for eusperm biased DEGs (B) and parasperm biased DEGs (C). Nodes represent GO subterms, with node size indicating the number of enriched genes. Nodes of the same color correspond to the same parent term, which is labeled in the figure. Edges indicate term similarity >30%, with thickness proportional to the similarity score. The figure was adapted from Metascape *(164)* output.

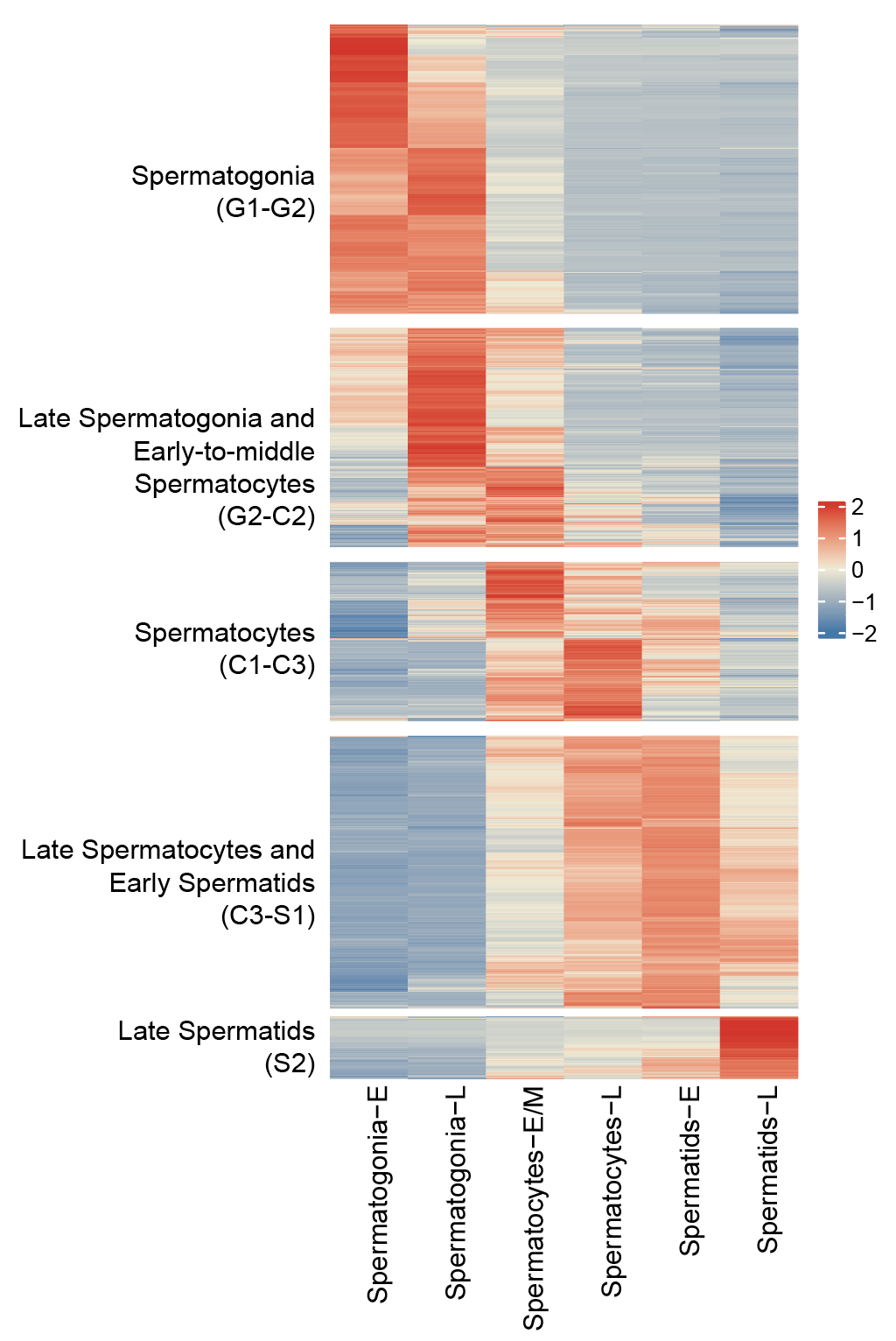

**Supplementary Fig. 24 Hierarchical clustering of expression of genes across different germ cell stages.** Heatmap showing the expression profiles of genes (rows) across different cell types (columns). Based on hierarchical clustering of average expression in each cell type, genes were grouped into 5 clusters: Spermatogonia; Late Spermatogonia and Early Spermatocytes; Spermatocytes; Late Spermatocytes and Early Spermatids; Late Spermatids. Color scale represents normalized expression levels from low to high.

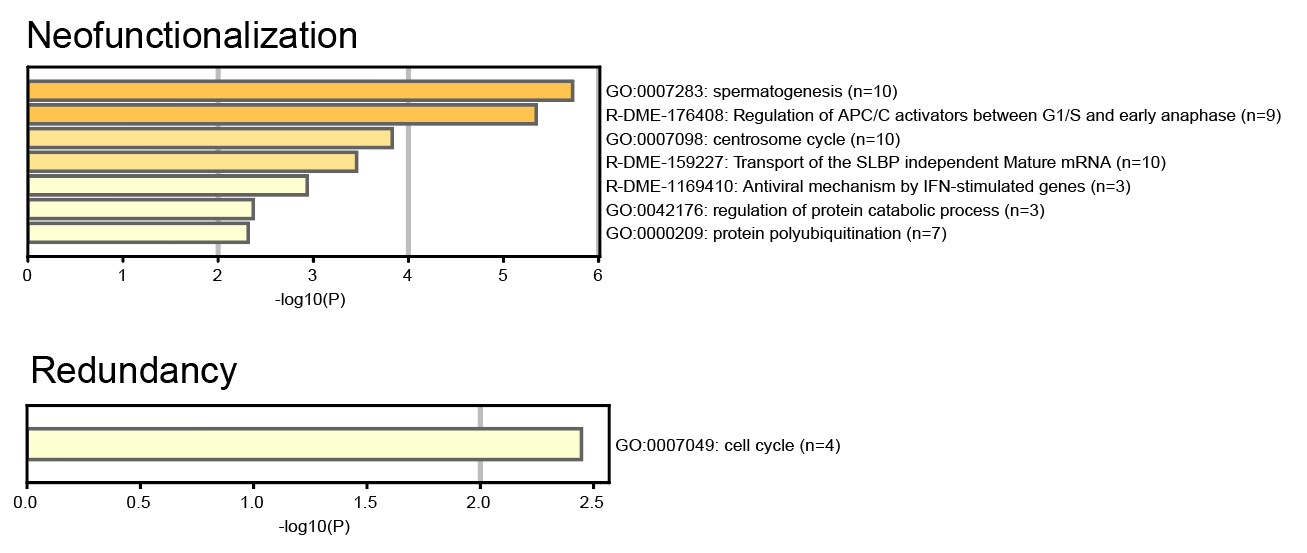

**Fig. S25 Gene Ontology (GO) enrichment of neofunctionalization genes and cell-type conserved genes.**

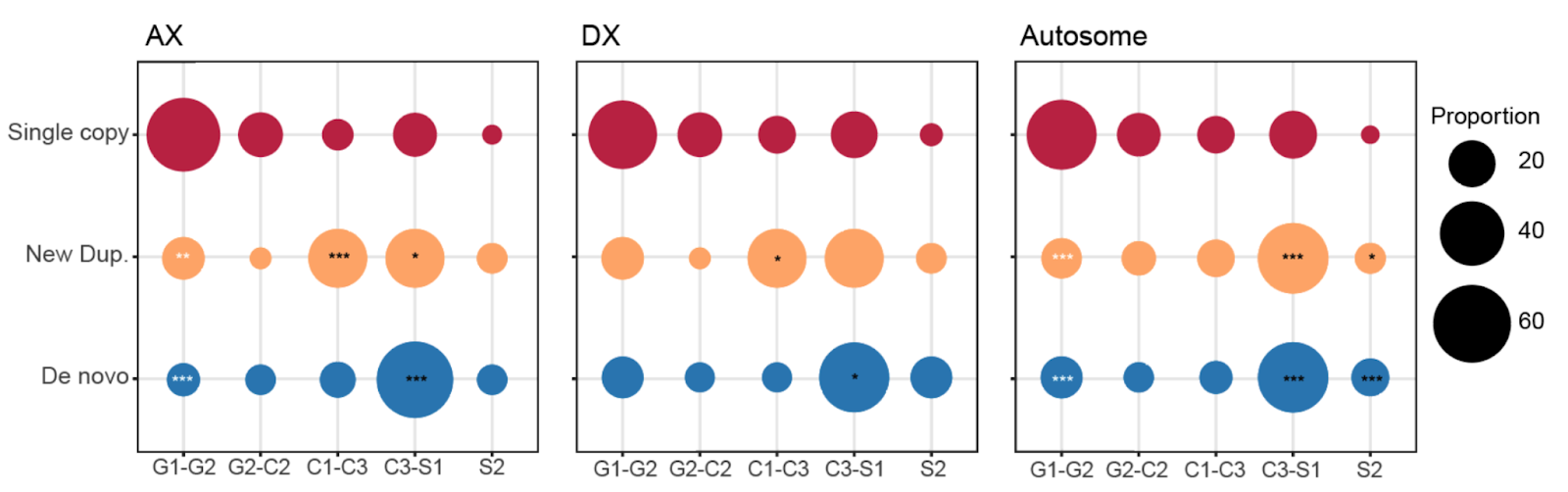

**Fig. S26 Distribution of predominantly expressed testis cell types of each category of genes on different chromosomes.** Asterisks denote a significant enrichment (black) or depletion (white) in newly duplicated genes or de novo genes compared with single-copy genes (one-sided Fisher’s exact test. *: *P* < 0.05, **: *P* <0.01, ***: *P* < 0.001).

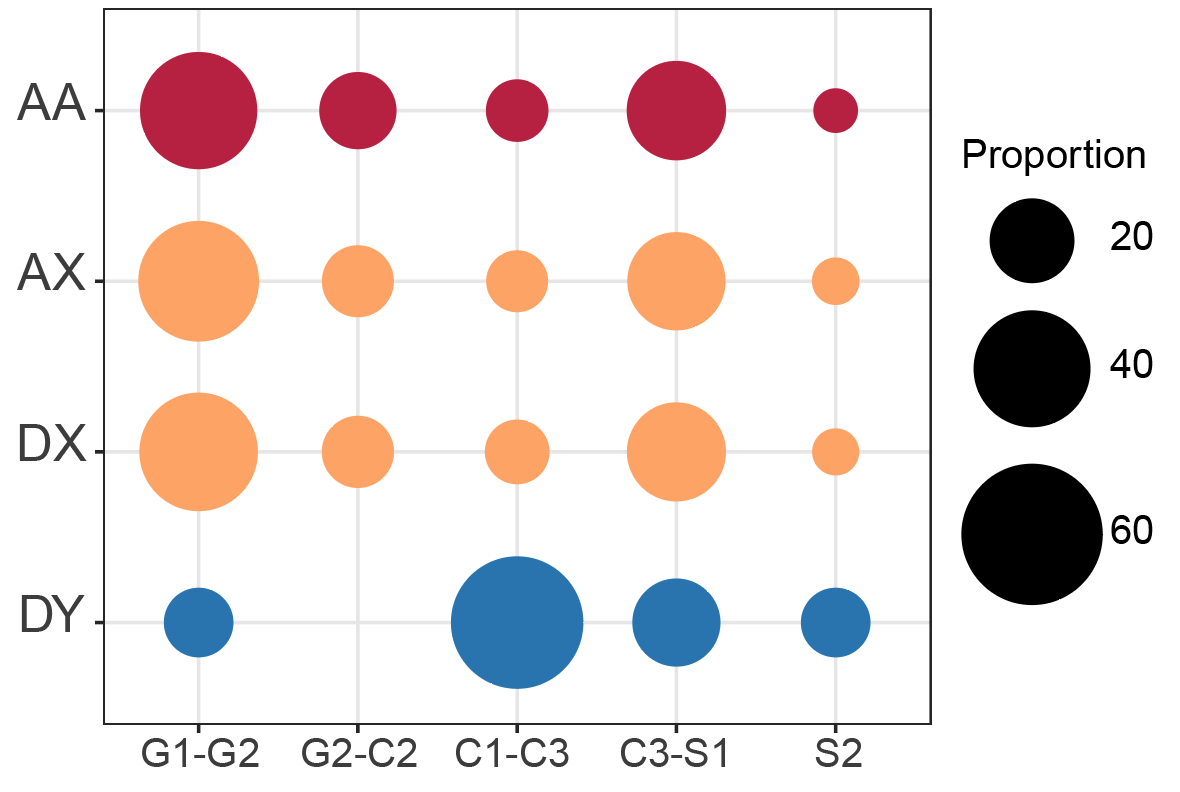

**Fig. S27 Distribution of predominantly expressed testis cell types of different chromosomes.**

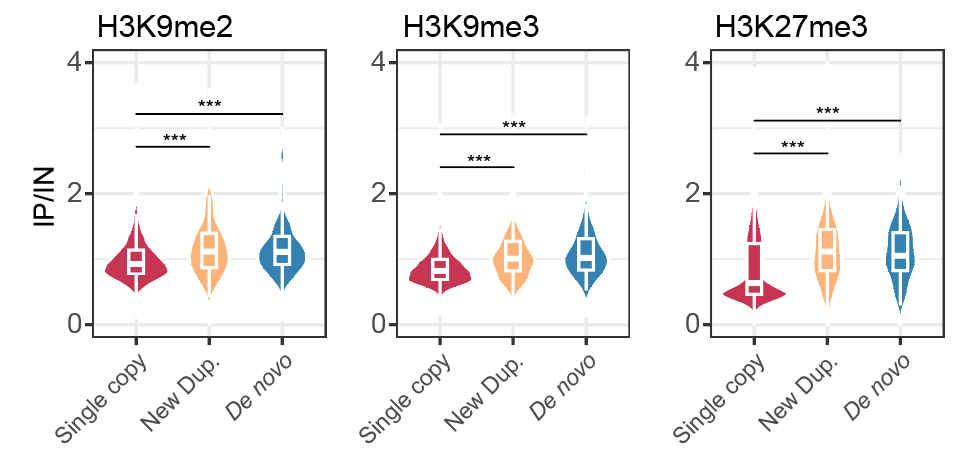

**Fig. S28 Comparisons of different histone modification enrichment between different gene categories.** ***: *P* < 0.001 Wilcoxon one-side
